# Longitudinal imaging of single cells in opaque tissue with cell chirp ultrasound

**DOI:** 10.64898/2026.10.06.756957

**Authors:** Rohit Nayak, Haein An, Trevor Fochtman, Zhiyang Jin, Mengtong Duan, Kaiwen Luo, Mikhail G. Shapiro

**Affiliations:** Division of Chemistry and Chemical Engineering, California Institute of Technology, Pasadena, California 91125, USA; Andrew and Peggy Cherng Department of Medical Engineering, California Institute of Technology, Pasadena, California 91125, USA; Division of Biology and Biological Engineering, California Institute of Technology, Pasadena, California 91125, USA; Howard Hughes Medical Institute, California Institute of Technology, Pasadena, California 91125, USA

## Abstract

Small numbers of cells can determine the fate of an entire organism. For example, metastatic cancer cells seed distant organs, therapy-resistant clones repopulate tumors, and T cells infiltrate and expand in tissues with matching antigens. Visualizing the activity of such cells in living organisms requires highly sensitive, specific and repeatable imaging in optically opaque tissues. Here we introduce *cell chirp*, an ultrasound imaging method for real-time imaging of single cells in living organs inaccessible to light. Cell chirp exploits the harmonic response of genetically encoded gas vesicles to frequency-coded ultrasound to sensitively and specifically visualize single cells in scattering background. We first develop this method in tissue-mimicking phantoms, demonstrating the ability to quantitatively detect single cells as validated by optical cell counts. We then establish the ability of cell chirp to image thousands of genetically labeled cells within a million-cell background tissue, track reporter gene expression over days, and suppress background signals by more than two orders of magnitude. Finally, we use cell chirp to track the fate of antigen-positive or antigen-negative tumor-cell minorities during weeks-long T-cell engager treatments. Cell chirp reveals the dynamics of sparsely distributed, genetically defined cells in opaque tissues.

## Introduction

Biologically consequential cell populations can be sparse—metastatic founders, resistant tumor clones, infiltrating immune cells and probiotic agents can appear as isolated cells or small clusters dispersed within much larger tissues. Imaging these rare cells would reveal when they appear, where they persist or expand, and when they disappear, connecting their early behavior to later tissue or organism outcomes. Tracking the dynamics of sparse cells requires longitudinal imaging with enough sensitivity to detect single cells and enough specificity to distinguish them from background tissue and imaging artifacts. Optical reporter genes such as fluorescent or luminescent proteins are unable to provide this performance in opaque tissues due to light scattering (**Fig. 1a**).

**Figure 1.**
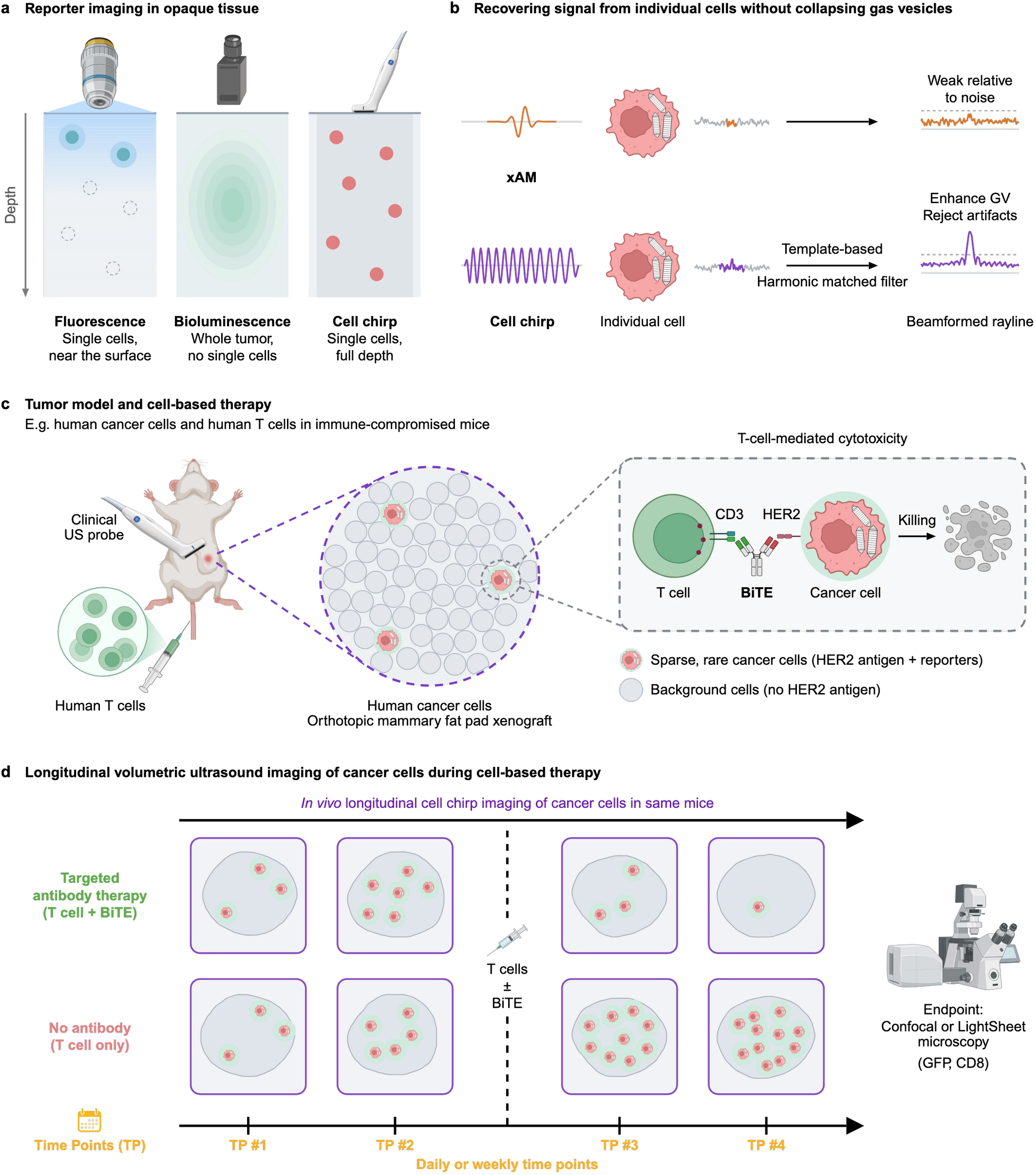
Frequency-encoded ultrasound enables longitudinal imaging of sparse reporter gene expression in opaque tissue. a, Reporter imaging in opaque tissue. Fluorescence can resolve individual cells near accessible surfaces, whereas whole-animal bioluminescence reports population-level signal without resolving cellular spatial organization at depth. Frequency-encoded ultrasound is designed to localize sparse reporter-expressing cells throughout intact tissue. b, Signal-recovery principle. Cell chirp extends excitation below the GV collapse threshold; amplitude modulation suppresses linear scattering, harmonic matched filtering concentrates the frequency-encoded nonlinear GV response and local background normalization rejects residual background. c, Population-specific tumor model. Reporter expression is restricted to a selected tumor-cell population in a heterogeneous tumor treated with human T cells with or without a HER2×CD3 BiTE. d, Longitudinal application. The same tumors are imaged at serial time points. Endpoint confocal imaging of GFP and CD8 provides an independent cellular readout of treatment response; light-sheet imaging of a separate cleared tumor compares three-dimensional reporter distributions with ultrasound. Created in BioRender. Shapiro, M. (2027) https://BioRender.com/idsurqg

Unlike light, ultrasound penetrates centimeters of living tissue, enabling large-scale volumetric imaging with high spatial (∼100 µm) and temporal (ms) resolution^19^. Mammalian acoustic reporter genes connect ultrasound signal to cells by encoding intracellular gas vesicles (GVs) – gas-filled protein nanostructures that scatter sound waves and produce nonlinear acoustic signals^1–4^. However, no ultrasound imaging method has been shown to reliably detect isolated cells against tissue background for real time and longitudinal imaging. Among existing imaging methods, BURST approaches single-cell sensitivity by irreversibly collapsing intracellular GVs, but in the process destroys the very agents that give the cells their acoustic contrast, limiting real-time and dynamic imaging^5^. Meanwhile, popular non-collapsing methods such as cross-wave amplitude modulation (xAM)^6^ and related AM approaches^7,14^ permit continuous readout, but fail to reliably detect single cells and reject artifacts from strongly-scattering interfaces.

Here, we introduce *cell chirp*, an ultrasound method that takes advantage of the unique acoustic properties of GVs to enable sensitive and specific real-time imaging of single cells. This method is based on two principles. The first is that GVs produce harmonic scattering by virtue of their reversible mechanical buckling under acoustic pressure, radiating sound waves at multiples of the transmitted ultrasound frequency. By modulating the transmit amplitude to be above and below GVs’ buckling threshold and collecting harmonics, it is possible to obtain highly specific signals from GV-expressing cells^14^. However, the signal thus obtained from single cells is too weak for reliable detection. To boost it, we can extend our transmitted pulse to a larger number of cycles, thereby increasing pulse energy – but this comes at the cost of spatial resolution, since in ultrasound this is based on time-of-flight. Such resolution loss can be overcome by using coded excitations, which allow received signals to be matched to an expected temporal pattern to compress them in time and, thereby, space. Chirps – sweeps of frequency within a transmitted pulse – are a common extended pulse pattern that has been used with microbubbles^8^.

In cell chirp, we combine amplitude modulation, chirp excitation and harmonic-filtered receive compression to recover enough signal to reliably detect single GV-expressing cells while greatly suppressing background scattering (**Fig. 1b**). By achieving this sensitivity while leaving the GVs intact, cell chirp allows continuous, dynamic and repeated imaging of sparse populations of genetically labeled mammalian cells. We implement cell chirp on a programmable ultrasound scanner using a clinical linear-array transducer. We demonstrate the ability of cell chirp to visualize single cells in phantoms, with detection counts in HEK-293T cells matching 99.8% of the optical cell-count estimate. In vivo, volumetric imaging of intact tumors enables mapping of the spatial distribution of a few thousand sparse GV-expressing cells across million-cell tumors, with cell chirp detecting more than 10 times as many reporter-associated contrast sources as xAM.

As a demonstration of longitudinal imaging, we use cell chirp to monitor genetically defined minority populations of tumor cells in orthotopic human tumor xenografts during treatment with primary T cells and a bispecific T-cell engager (BiTE) (**Fig. 1c**). By genetically labeling and imaging either antigen-positive or antigen-negative cells in separate experiments, we can use ultrasound to visualize direct and bystander treatment responses, while observing the absence of response in BiTE-free controls (**Fig. 1d**). We corroborate our non-invasive ultrasound observations with endpoint confocal microscopy. By enabling longitudinal, single-cell-sensitive readout of reporter gene expression, cell chirp provides unprecedented access to the dynamics of cells inside intact opaque tissues.

## Results

### Cell chirp enables sensitive, specific and repeated detection of sparsely distributed single cells

GVs exhibit three pressure-dependent scattering regimes: linear scattering below the buckling threshold, nonlinear scattering between the buckling and collapse thresholds, and irreversible collapse above the collapse threshold (**Fig. 2a**). Repeated detection of sparse reporter-expressing cells requires operating between these two thresholds, where reversible GV buckling generates nonlinear echoes without destroying the reporter. This places three demands on the measurement: sufficient sensitivity to detect isolated reporter sources, sufficient specificity to distinguish them from tissue background, and preservation of intact GVs for repeated imaging. To accomplish these goals, we combined amplitude modulation, extended chirp excitation, harmonic matched filtering and local background normalization in a single detection pipeline (**Fig. 1b** and **Fig. 2a**).

**Figure 2.**
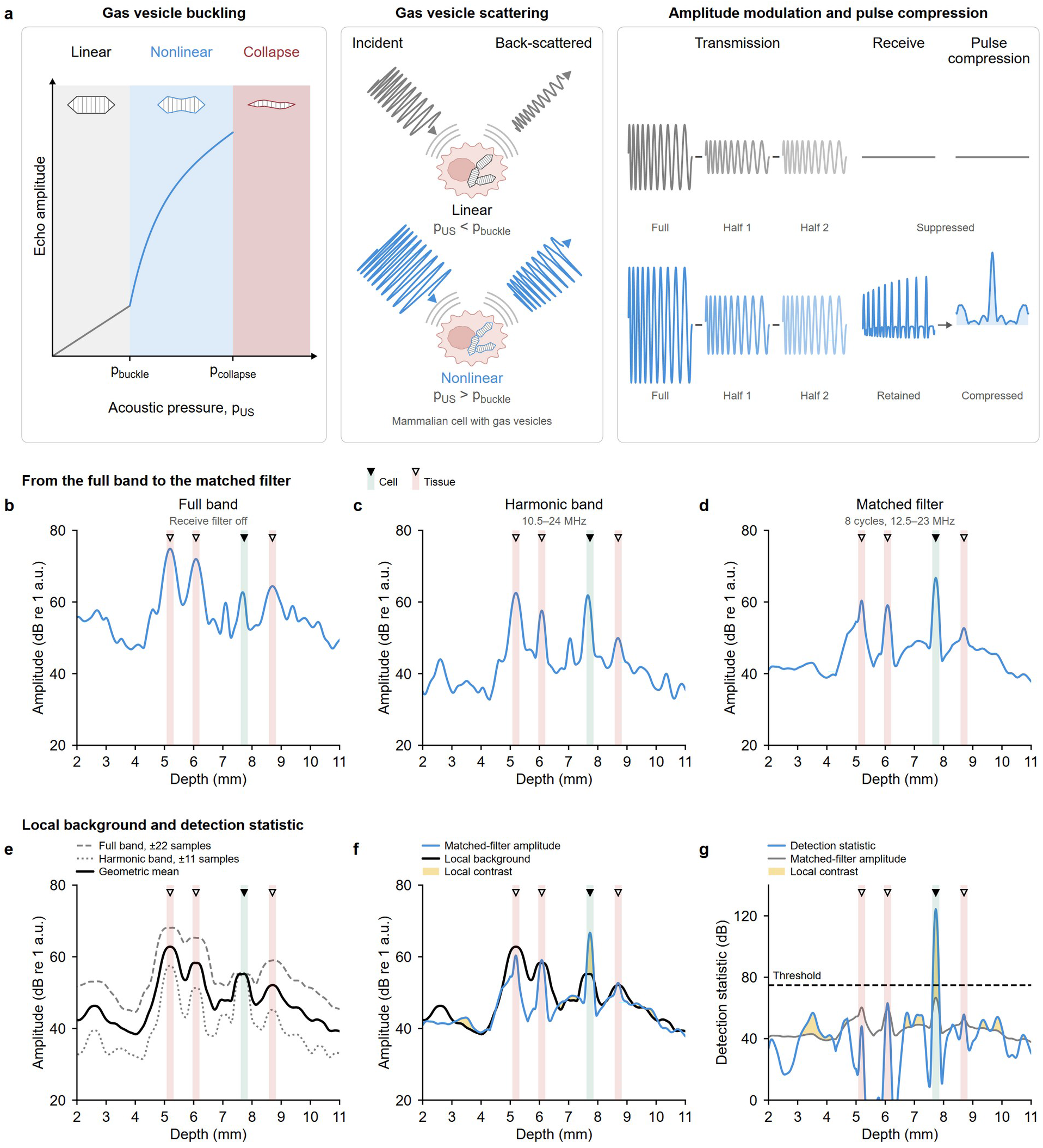
Frequency encoding recovers nonlinear GV echoes while rejecting tissue background. a, Pressure-dependent GV behavior and frequency-encoded signal recovery. GVs scatter linearly below the buckling threshold, buckle reversibly and generate nonlinear echoes between the buckling and collapse thresholds, and collapse irreversibly above the collapse threshold. Cell chirp imaging operates in the reversible nonlinear regime. One full-amplitude and two half-amplitude transmissions suppress the linear component by amplitude modulation while retaining nonlinear scattering; harmonic matched filtering then compresses the encoded response into a localized peak. b–g, Illustrative axial profiles showing the processing sequence for a reporter-associated echo among competing background echoes. The green band and filled triangle mark the reporter-associated response; red bands and open triangles mark background responses. b, Full-band envelope with the receive filter disabled. c, Harmonic-band envelope after 10.5–24-MHz filtering. d, Pulse-compressed envelope obtained with an eight-cycle, 12.5–23-MHz matched filter. e, Local background estimates from the full-band (±22 samples) and harmonic-band (±11 samples) responses and their geometric mean. f, Matched-filter amplitude relative to the local background; shading denotes local contrast. g, Detection statistic formed from the matched-filter response and local contrast, with the fixed threshold indicated. Panels b–f share a 20–80-dB amplitude scale referenced to 1 arbitrary unit. All waveforms and profiles are illustrative.

Chirp excitation extended the transmitted waveform over a 13-cycle frequency sweep from 10.5 to 7 MHz (**Fig. 2a**). The cell chirp lasted 1.76 µs, compared with 0.49 µs for the comparison xAM pulse. Hydrophone measurements showed little second-harmonic content in the transmitted cell chirp, indicating that the longer waveform did not introduce appreciable nonlinear transmission within the harmonic receive band (**Extended Data Fig. 1c,d**). Cell chirp and xAM were operated at 80% of their respective GV collapse-pressure thresholds (**Fig. 2a**).

In our cell chirp signal-recovery pipeline (**Fig. 2b–g and Extended Data Fig. 5f–k**), amplitude modulation suppresses linear scattering, while harmonic-band filtering and matched filtering isolate and compress the frequency-encoded nonlinear GV response into a localized peak (**Fig. 2, b–d**). Local estimates from the full-band and harmonic-band responses define the surrounding background (**Fig. 2e**). Normalization compares each peak with this background, and puncta are the peaks that exceed a fixed threshold (**Fig. 2f,g**).

We set the cell chirp threshold on uninduced, reporter-negative wells (79.6 dB) and used the previously described xAM analysis pipeline^14^ with a 40-dB envelope threshold (**Extended Data Fig. 4a,b** and Methods). Each threshold and its associated processing configuration remained fixed across all phantom datasets. Detection counts were defined by threshold crossings; parabolic fitting was applied only after detection to localize and display individual puncta (**Fig. 2g**, **Fig. 3g**, **Extended Data Fig. 3a,b** and Methods).

**Figure 3.**
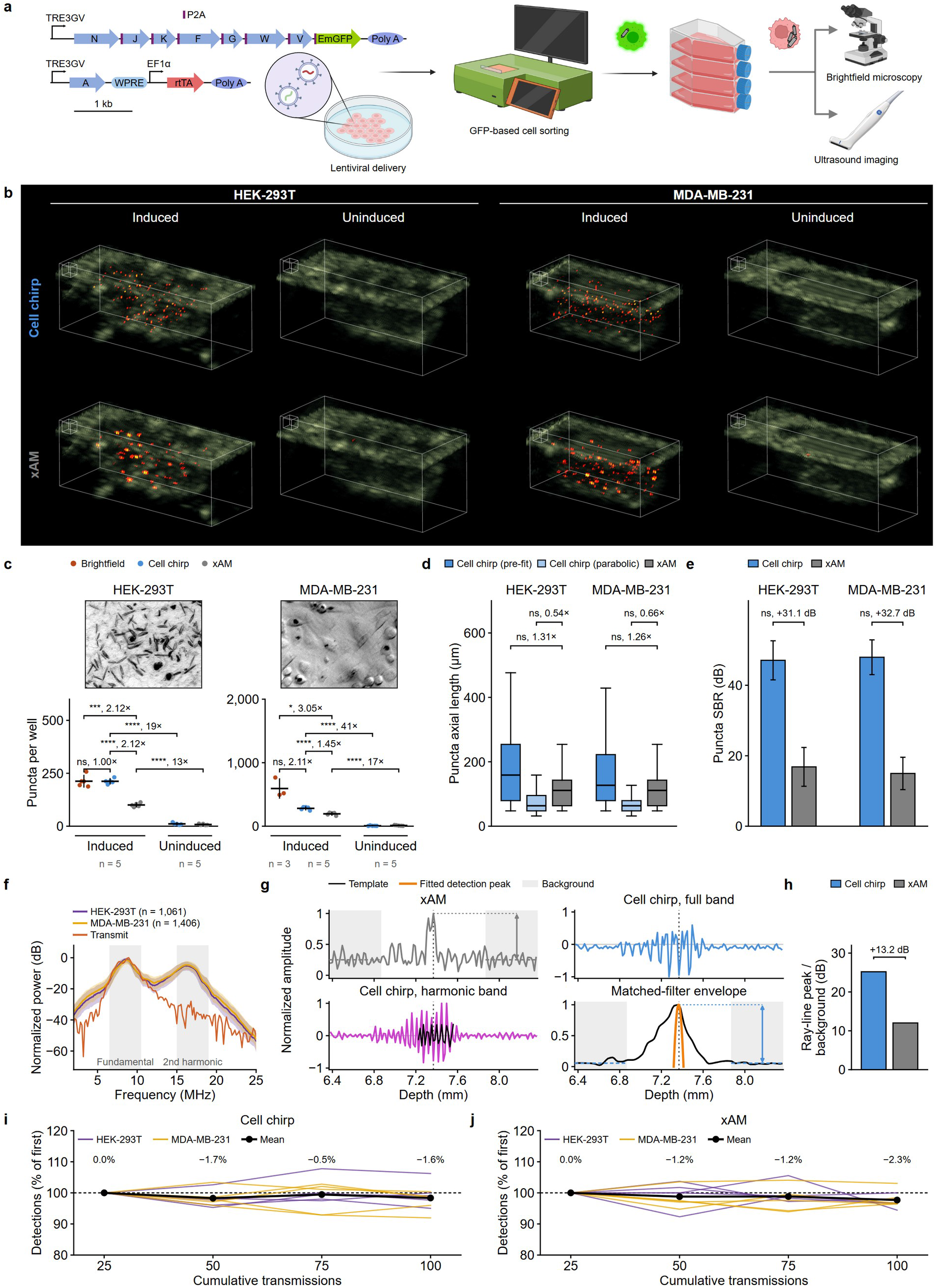
Cell chirp achieves single-cell-sensitive detection, recovers weak nonlinear reporter signals and supports repeated readout in phantoms. a, Generation and evaluation of inducible cellular acoustic reporters. The schematic shows lentiviral reporter delivery in MDA-MB-231 cells; HEK-293T cells used lentiviral rtTA delivery and PiggyBac reporter integration. Both lines were enriched by GFP-based sorting and evaluated by brightfield microscopy and ultrasound (Methods). b, Representative three-dimensional cell chirp and xAM detections overlaid on B-mode volumes from induced and uninduced HEK-293T and MDA-MB-231 wells. Scale cubes, 0.5 mm. c, Optical estimates of GV-positive cells and acoustic detections per well. For HEK-293T preparations, the optical estimate was 212.6 ± 27.4 GV-positive cells per well, cell chirp detected 212.2 ± 12.3 puncta and xAM detected 100.4 ± 8.9 puncta. For MDA-MB-231 preparations, the corresponding values were 593.4 ± 150.6, 281.2 ± 23.6 and 194.8 ± 20.7 (mean ± s.d.). Acoustic measurements used five wells per condition and cell line; optical estimates used five HEK-293T fields and three MDA-MB-231 fields. d, Axial length of threshold-positive cell chirp responses before fitting, their extent after three-point parabolic fitting and the axial length of xAM puncta. Detection counts were determined before fitting. Pooled medians were 158.8, 63.5 and 111.2 µm for HEK-293T cells and 127.0, 63.5 and 111.2 µm for MDA-MB-231 cells, respectively. e, Signal-to-background ratio of individual puncta, calculated with the specified local reference for each sequence; the cell chirp–xAM differences were +31.1 dB for HEK-293T cells and +32.7 dB for MDA-MB-231 cells. f, Median received power spectra with interquartile ranges for detected cell chirp puncta and the hydrophone-measured transmit spectrum; shaded regions mark the fundamental and second-harmonic bands (n = 1,061 HEK-293T and 1,406 MDA-MB-231 puncta). g, Measured rayline from MDA-MB-231 well 1 (plane 31, rayline 38), showing the xAM envelope, cell chirp full-band signal, cell chirp harmonic-band signal with the template overlaid and the matched-filter envelope with the fitted detection peak. Shaded regions define the background windows. h, Rayline peak above background before detector application. The xAM peak was 12.0 dB above background and the cell chirp matched-filter peak was 25.2 dB above background, a +13.2-dB difference. i,j, Detection counts (de-duplicated puncta, as in c) during repeated cell chirp (i) and xAM (j) imaging of ten reporter-expressing phantom wells: five MDA-MB-231 and five HEK-293T preparations. Each acquisition averaged 25 transmissions; the four measurements therefore correspond to 25, 50, 75 and 100 cumulative transmissions. Counts for each well were normalized to its first measurement. Colored lines denote individual wells; black points and lines denote the mean across all ten wells; the dashed line marks the first-measurement value. Mean changes from the first measurement were 0.0%, −1.7%, −0.5% and −1.6% for cell chirp and 0.0%, −1.2%, −1.2% and −2.3% for xAM. Punctum distributions are descriptive; sequence comparisons used wells as the independent units (n = 5 per cell line). *P < 0.05, ***P < 0.001, ****P < 0.0001; ns, not significant.

We next benchmarked cell chirp sensitivity against optical estimates of GV-expressing cells and evaluated specificity using reporter-negative controls. We used xAM as the gold-standard comparator for nonlinear, non-destructive cellular imaging^4,6,14,18,20–25^. We chose Human HEK-293T cells as an established cell type for mammalian acoustic reporter testing, and also tested MDA-MB-231 breast cancer cells as a prelude to our downstream orthotopic tumor experiments^3,4^. Both lines expressed doxycycline-inducible GVs and a green fluorescent protein (GFP) co-reporter and were enriched by GFP-based sorting (**Fig. 3a**). HEK-293T wells contained 600 induced cells mixed with 12,000 uninduced cells, and MDA-MB-231 wells contained 4,500 induced cells mixed with 13,500 uninduced cells. Cell chirp and xAM acquisitions were interleaved in five induced and five control wells from each line (**Fig. 3b**). Complete paired volumes from all 20 wells are presented in **Extended Data Fig. 2a**, and the brightfield fields used for the optical estimates are presented in **Extended Data Fig. 2b**.

Representative three-dimensional cell chirp volumes showed discrete puncta distributed throughout both induced samples (**Fig. 3b**). xAM also detected signal in the induced wells, but recovered a much smaller number of puncta than cell chirp. Both sequences produced few detections in the corresponding uninduced controls. In HEK-293T preparations, cell chirp detected 212.2 ± 12.3 puncta per well, closely matching the optical estimate of 212.6 ± 27.4 GV-positive cells per well; xAM detected only 100.4 ± 8.9 puncta per well under the same conditions (mean ± s.d.; **Fig. 3c** and **Extended Data Fig. 2c**). The agreement between cell chirp and the independent optical estimate supports single-cell detection sensitivity in this preparation. In MDA-MB-231 preparations, cell chirp detected 281.2 ± 23.6 puncta per well, which was lower than the optical estimate of 593.4 ± 150.6 GV-positive cells per well (**Fig. 3c** and **Extended Data Fig. 2c**), presumably due to lower per-cell GV expression (**Fig. 3c**). xAM detected only 194.8 ± 20.7 puncta per well.

To localize each detected punctum along the beam axis, we fitted a parabola to the three samples at the peak of its response after thresholding (Methods). Before fitting, the median above-threshold axial length of the cell chirp response was 159 µm in HEK-293T preparations and 127 µm in MDA-MB-231 preparations. After threshold-based detection, a three-point parabolic fit localized each peak and represented it with median axial lengths of 64 µm in both lines, compared with 111 µm for xAM (**Fig. 3d**).

Signal-to-background ratios of individual puncta were 31.1 dB higher for cell chirp than for xAM in HEK-293T preparations and 32.7 dB higher in MDA-MB-231 preparations (**Fig. 3e**). The cell chirp pipeline is enabled by the prominent second-harmonic response of GV-expressing cells (**Fig. 3f**, **Extended Data Fig. 1c,d** and **Extended Data Fig. 4c**). A representative MDA-MB-231 rayline traces the signal-recovery sequence from the xAM and cell chirp echoes through harmonic filtering, matched filtering and parabolic fitting (**Fig. 3g**). Before detector application, the cell chirp matched-filter peak was 25.2 dB above its local background, compared with 12.0 dB for xAM, a 13.2-dB difference in the same rayline (**Fig. 3h**). The corresponding processing stages for an induced phantom and reporter-negative control are shown in **Extended Data Fig. 3a,b**.

To investigate why xAM missed a subset of the puncta detected by cell chirp, we divided cell chirp detections into puncta co-detected by xAM and puncta detected only by cell chirp. Co-detected puncta had higher signal-to-background ratios and stronger harmonic content than cell chirp puncta in both cell lines (**Extended Data Fig. 4d,e**). The additional cell chirp detections therefore carried weaker nonlinear GV signatures that xAM failed to recover.

Finally, we tested whether repeated imaging depleted the reporter response. Across four acquisitions spanning 100 transmissions, mean cell chirp detection counts remained within 2% of the first acquisition and fluctuated in both directions between scans; xAM counts changed similarly little over the same series (**Fig. 3i,j**). Cell chirp therefore combined single-cell sensitivity and reporter specificity with repeated detection without using irreversible GV collapse as the detection event.

### Cell chirp enables specific, longitudinal detection of sparse reporters in intact tumors

After establishing the basic capabilities of cell chirp in phantoms, we tested whether this performance translated to living tissue. As our testing ground, we selected orthotopic breast tumors, where nonlinear tissue echoes and interfacial reflections present a more challenging background. As the first test, we asked whether cell chirp could detect sparse GV-expressing cells in the tumor, follow the reporter-positive population through daily imaging in the same animal, and distinguish GV-dependent signals from pre-induction and post-collapse controls.

We generated orthotopic tumors by implanting a mixture of 20 thousand MDA-MB-231 cells carrying the doxycycline-inducible GV and GFP reporter and one million reporter-negative cells. We imaged 4 tumors before induction and at 3 subsequent daily timepoints, while inducing expression with systemic doxycycline (dox). We then collapsed the GVs and repeated the acquisition to determine which detections depended on intact reporters (**Fig. 4a,b**). Paired cell chirp and xAM volumes from every animal and time point are presented in **Extended Data Fig. 5a**. We set the in vivo cell chirp threshold on the four post-collapse tumor volumes (74.8 dB) and used the same 40-dB envelope threshold for xAM (Methods). Each threshold and its associated processing configuration remained fixed across the complete in vivo dataset (**Extended Data Fig. 5l,m**).

**Figure 4.**
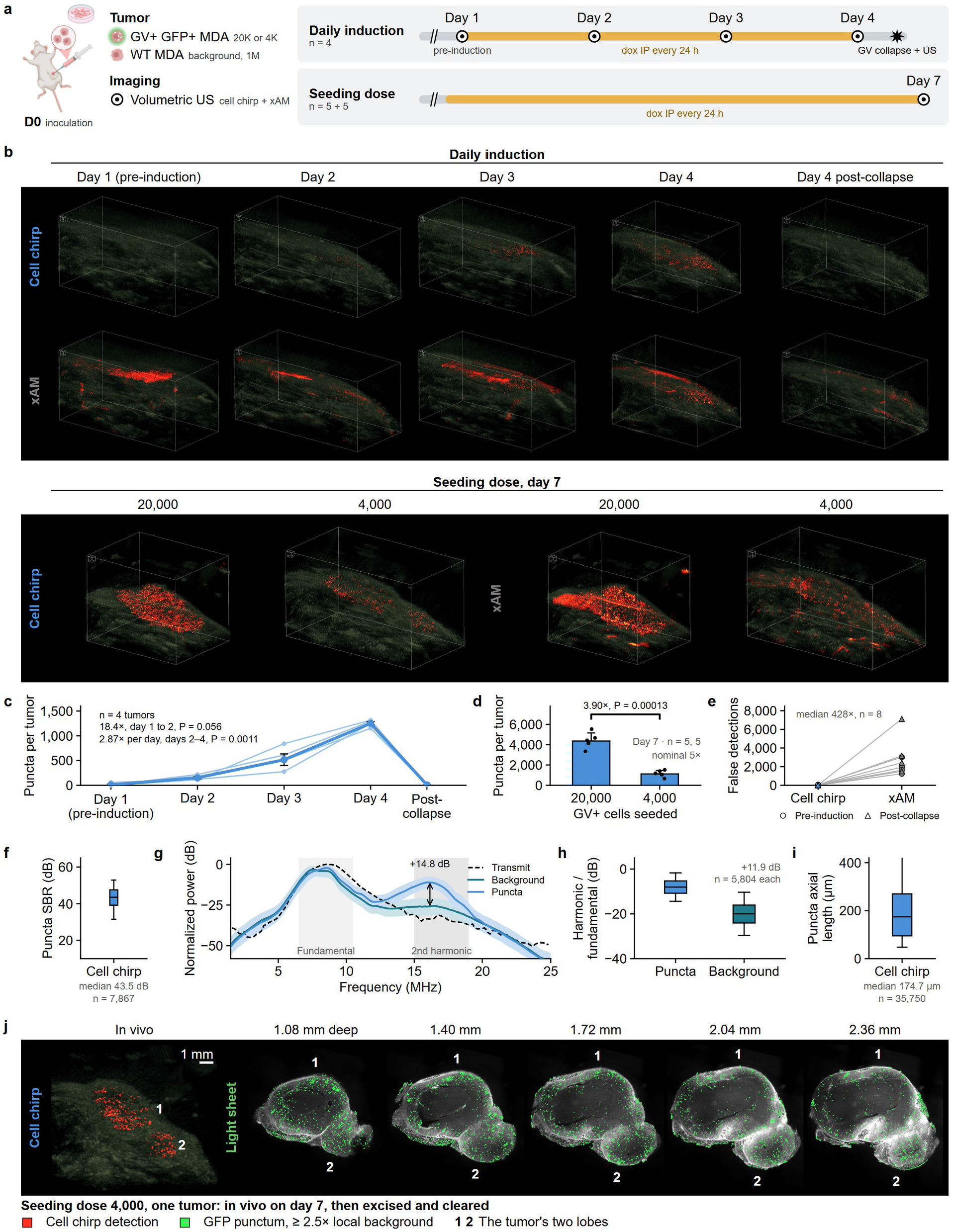
Cell chirp enables specific, longitudinal detection of sparse reporters in intact tumors. a, Experimental designs for testing reporter induction, longitudinal readout, GV dependence and sensitivity to reporter-cell input in vivo. Orthotopic mammary fat pad tumors contained either 2 × 10⁴ or 4 × 10S reporter-positive MDA-MB-231 cells among 1 × 10⁶ reporter-negative cells. Four tumors in the daily-induction study were imaged before induction, 24 h after each of three daily doxycycline doses and immediately after GV collapse. Separate cohorts initiated with 20,000 or 4,000 reporter cells were imaged on day 7 (n = 5 tumors per group). b, Representative volumetric cell chirp and xAM detections overlaid on B-mode. The upper rows follow one tumor through induction and collapse; the lower row compares tumors initiated with 20,000 or 4,000 reporter cells. Scale cubes, 0.5 mm. Complete paired volumes are presented in **Extended Data Fig. 5.** c, Cell chirp detections in the same four tumors across induction. Thin lines denote individual tumors and the thick line denotes the geometric mean: 8 puncta before induction, 152 after the first dose, 516 after the second and 1,251 after the third, followed by 9 after collapse. The first 24-h increase was 18.4-fold (P = 0.056; two-sided one-sample t-test on log ratios, n = 4 tumors). Collapse removed 99.3% of pooled induced detections. d, Day-7 detections from tumors initiated with 20,000 or 4,000 reporter cells. The 20,000-cell group produced 3.90-fold more puncta (P = 0.00013; two-sided Welch t-test on log counts, n = 5 tumors per group). e, Paired false-detection counts for cell chirp and xAM in the eight scans carrying no induced reporter signal: four pre-induction and four post-collapse volumes. Cell chirp reduced false detections by a median factor of 428 relative to xAM (range, 51.2–1,242). f, Signal-to-background ratio of each in vivo cell chirp punctum relative to the background of the same counting region after GV collapse (median, 43.5 dB; n = 7,867 puncta). g, Median spectra of detected puncta and depth-matched background from the same planes, with the transmit spectrum overlaid (n = 5,804 per class). At 16.15 MHz, the received spectral power at cell chirp-detected puncta exceeded that of depth-matched background by 14.8 dB; the mean difference across 15–19 MHz was 11.03 dB. h, Ratio of harmonic-band to fundamental-band power for the same puncta and background samples; the median separation was 11.9 dB. i, Axial length of in vivo cell chirp detections before fitting (median, 174.7 µm; n = 35,750 detections). Boxes in f, h and i show the median and interquartile range. j, Same-tumor comparison after day-7 imaging of a tumor initiated with 4,000 reporter cells (n = 1 tumor). Left, in vivo cell chirp detections. Right, endogenous GFP fluorescence in five optical planes acquired after tumor excision, clearing and light-sheet microscopy at the indicated depths. GFP puncta were displayed at least 2.5-fold above local background. Numerals 1 and 2 identify the two tumor lobes. All images are shown at the same physical scale; scale bar, 1 mm. The light-sheet planes were reoriented for display to align gross lobe anatomy with the cell chirp rendering; the modalities were not pointwise registered.

Before induction, cell chirp produced few detections within the tumor-containing B-mode volume. Discrete puncta appeared 24 hours after induction, became progressively more numerous over the next 3 days, and largely disappeared after GV collapse (**Fig. 4b**). Across subjects, the geometric mean number of puncta detected by cell chirp increased from 8 before induction to 1,251 on day 4 (**Fig. 4c**). GV collapse eliminated 99.3% of the induced detections, returning the geometric mean to 9 puncta per tumor (**Fig. 4c** and **Extended Data Fig. 5a,c–e**).

Compared to cell chirp, xAM produced many more artifactual detections before GV induction and after collapse, making the induction-dependent reporter signal difficult to distinguish on images (**Fig. 4b** and **Extended Data Fig. 5a**). At fixed detection thresholds, cell chirp reduced false detections by a median factor of 428 relative to xAM (**Fig. 4e**). When the xAM threshold was raised to give the same false-detection rate as cell chirp, cell chirp detected 27.6 times as many puncta as xAM in the induced tumors (**Extended Data Fig. 5n,o**).

In vivo signals retained the acoustic characteristics established in phantoms. The median signal-to-background ratio of in vivo puncta was 43.5 dB, relative to the background of the same counting region after GV collapse (**Fig. 4f**). In the received spectra, puncta carried more power than depth-matched background across the second-harmonic band, by 11.03 dB on average and by up to 14.8 dB (**Fig. 4g**). Harmonic-to-fundamental power separated puncta from background by a median of 11.9 dB (**Fig. 4h**). Before parabolic fitting, the median axial length of detections was 175 µm in tumors and 127 µm in MDA-MB-231 phantoms (**Figs. 3d and 4i**). These measurements established that cell chirp provides exceptional ability to generate nonlinear spectral contrast, provide precise axial localization, and separate positive signal from tissue background in vivo.

To assess ability of cell chirp to capture relative quantities of cells in tissue, we reduced the reporter-positive cell input from 20,000 to 4,000 cells among 1M reporter-negative cells. Cell chirp images contained correspondingly fewer puncta (**Fig. 4b** and **Extended Data Fig. 5b**). On day 7, tumors initiated with 20,000 reporter cells produced 3.90 times as many puncta as tumors initiated with 4,000 reporter cells (geometric means; 4,416 ± 800 versus 1,161 ± 332 puncta, mean ± s.d.; **Fig. 4d**). The relatively close correspondence between the 5x difference in input cells and the 4x difference in detected signal is remarkable given the variability typically associated with visualizing relatively large tissue volumes.

To compare the general spatial distribution of reporter-positive cells detected by cell chirp to post-mortem optical microscopy, we excised and cleared a tumor after day-7 ultrasound imaging and examined co-expressed GFP by volumetric light-sheet microscopy (**Fig. 4j**). Both imaging modes revealed two tumor lobes with sparse puncta distributed across both lobes. While puncta-wise registration is outside the scope of this study, the qualitative distribution was similar between in vivo cell chirp and post-mortem optical imaging.

Having established the basic in vivo performance of cell chirp, we next evaluated its ability to reveal the dynamics of a rare cell population during cell therapy.

### Cell chirp resolved a population-specific response to antigen-targeted therapy

While antigen-targeted therapies are highly effective in eliminating cancer cells^9^, tumor heterogeneity can leave cellular subpopulations within the tumor intact, leading to treatment resistance or recurrence^10^. Bulk measurements of tumor growth are unable to capture the internal population dynamics of such treatments, obscuring insights that could otherwise be useful for understanding cancer biology or developing improved treatments. We asked whether cell chirp could make these otherwise hidden dynamics observable by following genetically defined minority cell populations within tumors undergoing antigen-targeted treatment.

For our first set of experiments, we established orthotopic human tumors containing reporter-positive and HER2-positive human MDA-MB-231 cells as a minority among HER2-negative, reporter-negative WT MDA-MB-231 cells. We treated the tumors with adoptively transferred human peripheral blood mononuclear cell (PBMC)-derived T cells and runimotamab, a HER2×CD3 BiTE (**Fig. 5a**). Under these conditions, the BiTE is expected to target the T cells toward the reporter-positive cells, leading to their elimination.

**Figure 5.**
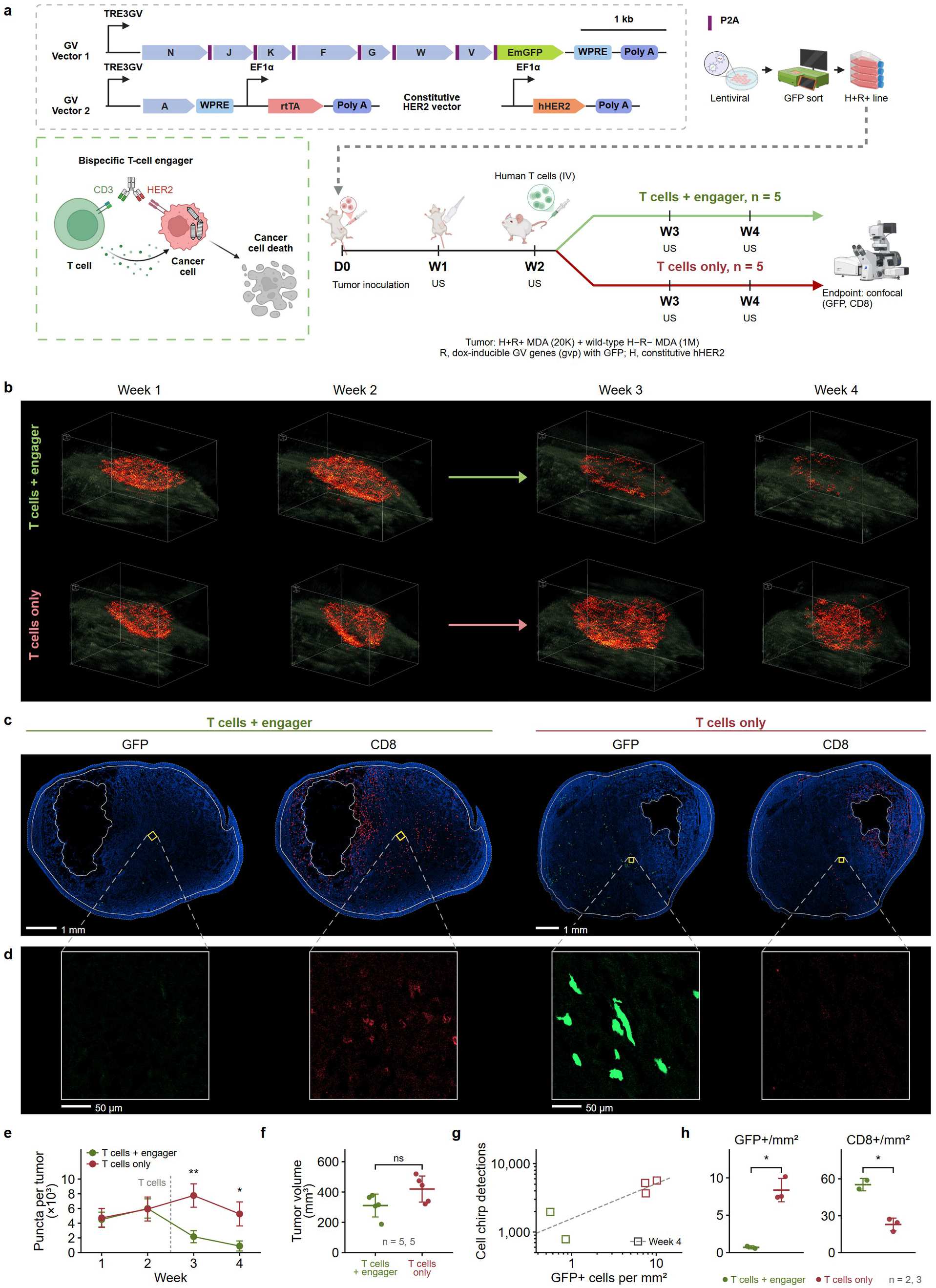
Cell chirp reveals a treatment response in an antigen-targeted tumor-cell population that is obscured by tumor volume. a, Human effector–target model and study design. Orthotopic tumors contained 2 × 10⁴ reporter-positive, HER2-positive MDA-MB-231 cells among 1 × 10⁶ reporter-negative parental MDA-MB-231 cells without ectopic HER2 expression. All mice received human PBMC-derived T cells after week 2; one group also received the HER2×CD3 BiTE runimotamab every 48 h (n = 5 mice per group). The same tumors were imaged twice before and twice after treatment. b, Longitudinal cell chirp volumes from one representative mouse per group. Red puncta mark the HER2-positive reporter population; complete cell chirp volumes are shown in **Extended Data Fig. 6**. Scale cubes, 0.5 mm. c, Endpoint whole-tumor confocal sections from the representative mice, showing GFP-positive reporter cells and CD8-positive T cells. Scale bars, 1 mm. d, Raw 250-µm fields from the positions marked in c. Scale bars, 50 µm. e, Longitudinal reporter detections, mean ± s.d. Counts were comparable before treatment and were 3.85-fold lower at week 3 and 8.42-fold lower at week 4 in the T cells + BiTE group than in T-cell-only controls (P = 0.0028 and P = 0.016, respectively; n = 5 mice per group). f, Endpoint tumor volume, mean ± s.d. The T cells + BiTE group was 0.74 times the T-cell-only control volume (P = 0.066; n = 5 mice per group). g, Week-4 cell chirp detections plotted against endpoint GFP-positive cell density on log scales for animals with microscopy measurements (r = 0.86, P = 0.064). h, Endpoint GFP-positive and CD8-positive cell densities, shown as per-mouse values with mean ± s.d. GFP-positive cell density with BiTE was 0.08 times that in T-cell-only controls (P = 0.022), whereas CD8-positive cell density was 2.14-fold higher (P = 0.013). One section per animal was analyzed: n = 2 T cells + BiTE and n = 3 T cells only for GFP, and n = 2 per group for CD8. Weekly and endpoint comparisons used two-sided Welch t-tests with mice as the independent units; count tests used log_10_-transformed values and fold differences are ratios of geometric means. *P < 0.05, **P < 0.01; ns, not significant.

We seeded the tumors with 20,000 HER2+ reporter cells and 1M HER2–, reporter-negative cells and imaged them over 4 weeks. All animals received human T cells after 2 weeks, and a subset subsequently received BiTE treatment every 48 h. Tumors were imaged twice before and twice after treatment initiation (**Extended Data Fig. 6**). Before treatment, all tumors showed similar spatial distributions of HER2+ reporter cells. After BiTE administration, ultrasound-detectable puncta diminished in the treated tumor while remaining abundant in the T-cell-only control (**Fig. 5b**). Across the cohort, BiTE-treated tumors contained 3.85-fold fewer reporter puncta at week 3 and 8.42-fold fewer at week 4 relative to T-cell-only controls (P = 0.0028 and P = 0.016, respectively; **Fig. 5e** and **Extended Data Fig. 6**). Meanwhile, overall tumor volume between the two groups only differed by 26% (P = 0.066; **Fig. 5e,f**), demonstrating the additional information provided by imaging a specific, rare cell sub-population relative to bulk tissue.

Endpoint confocal microscopy confirmed the population-specific response measured by cell chirp. GFP-positive cell density was lower in BiTE-treated animals than in T-cell-only controls (treated-to-control ratio, 0.08; P = 0.022; **Fig. 5c,d,h** and **Extended Data Fig. 7**). CD8-positive T-cell density was also 2.14-fold higher after BiTE treatment (P = 0.013; **Fig. 5h**).

These experiments established that cell chirp could distinguish the longitudinal response of an antigen-defined minority from the composite response of a heterogeneous tumor.

### Cell chirp reveals a bystander response in sparse antigen-negative tumor cells

Having validated the ability of cell chirp imaging to follow the elimination of a rare antigen-positive cell population by a matching targeted immunotherapy, we tested the ability of this imaging method to answer a less-settled question: what happens to rare antigen-negative cells when surrounding antigen-positive cells are being killed by immune cells? Will they survive and proliferate, or will they also come under immune attack as bystanders^12,13^? To study this question in orthotopic breast tumor immunotherapy, we reversed the antigen and reporter configuration from the preceding experiment, expressing GVs in HER2– cells, while leaving HER2+ cells without an ultrasound reporter.

We established orthotopic tumors containing 20,000 reporter-positive MDA-MB-231 cells without ectopic HER2 expression and 1M HER2-positive, reporter-negative cells (**Fig. 6a**). As previously, we treated animals with the HER2-BiTE-mediated primary human T-cell therapy, with a subset of the mice receiving the T cells alone without the BiTE. We imaged the tumor weekly, twice before and twice after treatment (**Fig. 6b**, **Extended Data Fig. 8**).

**Figure 6.**
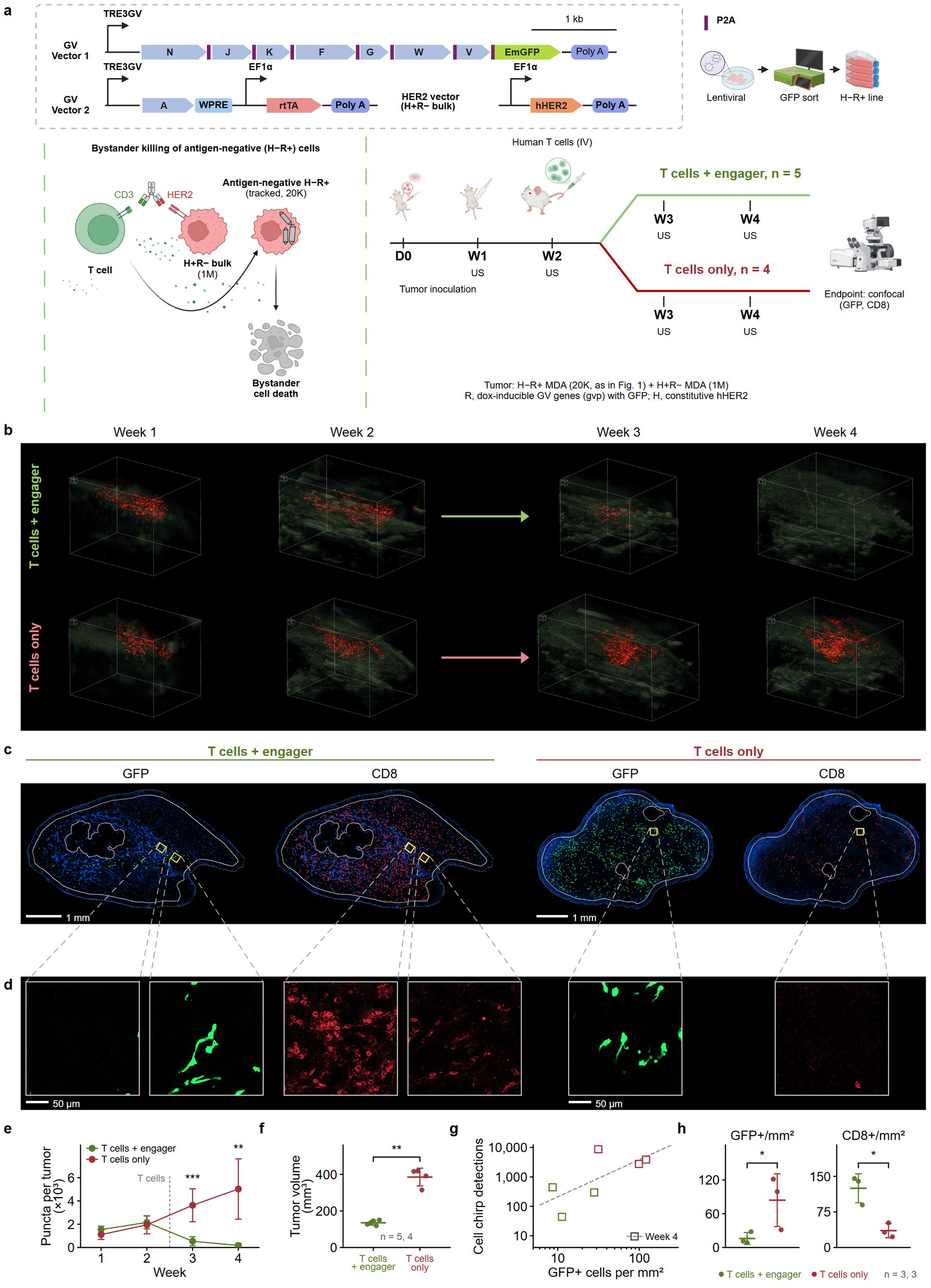
Cell chirp reveals a longitudinal treatment response in sparse antigen-negative tumor cells. a, Reciprocal reporter assignment and study design. Orthotopic tumors contained 2 × 10⁴ reporter-positive MDA-MB-231 cells without ectopic HER2 expression among 1 × 10⁶ HER2-positive, reporter-negative MDA-MB-231 cells. The BiTE was therefore directed at the surrounding HER2-positive population rather than the imaged cells. All mice received human T cells after week 2; one group also received the HER2×CD3 BiTE every 48 h (n = 5 mice with T cells + BiTE and n = 4 with T cells only). b, Longitudinal cell chirp volumes from one representative mouse per group. Red puncta mark the HER2-negative reporter population; complete cell chirp volumes are shown in **Extended Data Fig. 8**. Scale cubes, 0.5 mm. c, Endpoint whole-tumor confocal sections from the representative mice, showing GFP-positive reporter cells and CD8-positive T cells. Scale bars, 1 mm. d, Raw 250-µm fields from the positions marked in c. Scale bars, 50 µm. e, Longitudinal reporter detections, mean ± s.d. Counts fell after treatment in all five BiTE-treated tumors and remained 69 –99% below each tumor’s week-2 value at week 4, whereas counts increased in all four T-cell-only controls. Between groups, counts were 7.6-fold lower at week 3 and 51.3-fold lower at week 4 with BiTE (P = 0.0007 and P = 0.0018, respectively). f, Endpoint tumor volume, mean ± s.d.; the T cells + BiTE group was 0.35 times the T-cell-only control volume (P = 0.0013). g, Week-4 cell chirp detections plotted against endpoint GFP-positive cell density on log scales for animals with microscopy measurements (r = 0.69, P = 0.127). h, Endpoint GFP-positive and CD8-positive cell densities, shown as per-mouse values with mean ± s.d. GFP-positive cell density was 16.0 ± 10.5 cells mm⁻² with T cells + BiTE and 84.1 ± 46.9 cells mm⁻² with T cells alone (0.19-fold, P = 0.042). CD8-positive cell density was 3.64-fold higher with BiTE (P = 0.016; n = 3 mice per group). Weekly and endpoint comparisons used two-sided Welch t-tests with mice as the independent units; count tests used log_10_-transformed values and fold differences are ratios of geometric means. *P < 0.05, **P < 0.01, ***P < 0.001; ns, not significant.

Our longitudinal series of cell chirp images showed the HER2-negative cell population declining after BiTE treatment while remaining abundant in the T-cell-only tumor (**Fig. 6b, e** and **Extended Data Fig. 8**). Relative to immediately before treatment, punctum counts decreased in all five BiTE-treated subjects and remained below pretreatment values at the endpoints, declining by 69–99% (**Fig. 6e** and **Extended Data Fig. 8a**). BiTE-treated tumors contained 7.6-fold fewer puncta than T cell-only controls at week 3 and 51.3-fold fewer at week 4 (P = 0.0007 and P = 0.0018, respectively; **Fig. 6e** and **Extended Data Fig. 8**). Meanwhile, the overall tumor shrank by 65% relative to T-cell-only controls, indicating successful killing of the background HER2+ majority (P = 0.0013; **Fig. 6f**).

Endpoint whole-tumor confocal microscopy corroborated the reduction of GFP-positive bystander cells in BiTE-treated tumors relative to T cell-only controls (**Fig. 6b–d, h** and **Extended Data Fig. 9**). BiTE treatment increased intratumoral CD8-positive cell density 3.64-fold (P = 0.016; n = 3 tumors per group; **Fig. 6h**). Thus, the HER2-negative reporter population declined within a CD8-infiltrated tumor microenvironment.

On an individual-tumor basis, tumors with more cell chirp puncta at week 4 contained more GFP-positive cells under optical microscopy after adjustment for tumor model (r = 0.65, P = 0.040; n = 11 animals; **Figs. 5g and 6g**), suggesting that ultrasound imaging quantitatively reflects the relative abundance of rare cells in living tissue.

Together, these results show that cell chirp imaging is capable of revealing both expected and unexpected sparse cell population dynamics in living opaque tissues.

## Discussion

Our results establish cell chirp imaging as a sensitive and highly specific method for repeatedly visualizing sparse cellular populations in optically opaque tissues. In phantoms, the detection of puncta validated by optical cell counts supports single-cell sensitivity. In vivo, restricting reporter expression to genetically defined cell populations allowed us to measure the treatment responses of HER2-positive and HER2-negative minorities during targeted immunotherapy. Together, these findings show that cell chirp enables longitudinal measurement of selected cell populations whose treatment responses are not evident from bulk tumor size^9,10^.

Compared with conventional xAM, cell chirp improved both background rejection (by 428-fold) and reporter recovery (by 27.6-fold when compared at the same false-detection rate). Harmonic matched filtering compressed the frequency-encoded nonlinear response, while local normalization separated reporter peaks from surviving background. The combined gain in weak-signal recovery and background rejection allows cell chirp to provide robust detection of sparse cell distributions practical in deep intact tissue (>1 cm), where missed signals and false detections would otherwise confound the apparent population response.

The treatment experiments demonstrate the value of measuring selected minorities during a biological perturbation. Labeling HER2-positive cells revealed a pronounced response in the antigen-targeted population, with 3.85-fold and 8.42-fold fewer puncta at weeks 3 and 4, whereas the endpoint difference in whole-tumor volume was not statistically significant. In a reciprocal experiment, reversing the reporter and antigen configuration addressed whether cells without ectopic HER2 expression also responded to treatment as bystanders, providing a clear affirmative answer. Endpoint microscopy corroborated the lower abundance of GFP-positive cells in BiTE-treated tumors. These findings are consistent with previous studies showing that BiTEs can produce bystander killing, including cytokine-mediated effects on HER2-negative cells after activation at HER2-positive targets^12,13^. Our measurements support a substantial bystander contribution in the MDA-MB-231 model and provide a way to investigate how antigen heterogeneity shapes solid-tumor composition and treatment response over time.

Beyond the examples shown in this initial study, the ability to repeatedly localize genetically selected reporter populations in intact tissue could support longitudinal studies of engineered cell therapies^15^, residual treatment-resistant tumor cells and early metastatic dissemination. Cell chirp complements optical approaches rather than replacing them: bioluminescence provides population monitoring without effective spatial resolution, while intravital microscopy provides detailed cellular information within a restricted field of view^16,17^. Cell chirp adds repeated, noninvasive volumetric localization of cells that can be spatially registered to tumor boundaries, vascular anatomy, internal echotexture and cystic or necrotic regions.

Further development should connect longitudinal punctum measurements to tumor anatomy and growth. The current study lacked a volumetric anatomical reference for registering tumors across time, preventing direct comparison of population-specific changes with local tumor growth kinetics. Co-registering cell chirp detections with serial three-dimensional ultrasound anatomy would allow sites of persistence, regression, and recurrence to be followed within the same anatomical framework. Cell chirp’s punctum-based readout is most informative for sparse populations; as reporter signals overlap at higher densities, discrete counting becomes less informative, and regional xAM or HxAM measurements may provide a complementary readout.^6,14^ Broader use will require reporter-engineering strategies that achieve consistent expression across cell types and within each cell population, together with validation in varied tissue backgrounds.

Cell chirp enables sensitive, highly specific and repeated imaging of selected cellular populations in intact tissue without relying on irreversible GV collapse. This capability should enable an upswing in understanding of how biologically consequential cell populations respond to and drive the behavior of whole tissues and organisms.

## Methods

### Cell lines and culture conditions

HEK-293T cells and human MDA-MB-231 breast adenocarcinoma cells were maintained at 37 °C in a humidified incubator with 5% CO₂. Complete growth medium was prepared by supplementing 500 ml 1× DMEM with 50 ml fetal bovine serum, 5 ml penicillin–streptomycin, 10 ml 1 M HEPES and 5 ml 100× non-essential amino acids. Cultures were routinely passaged before reaching confluence. MDA-MB-231 cells were used for the in vitro phantom, orthotopic tumor and treatment studies; HEK-293T cells provided an established reporter-engineering and optical–acoustic benchmarking background.^3,4^

### Lentiviral vector architecture

The inducible mammalian acoustic reporter program in MDA-MB-231 cells comprised a TRE3G-driven structural-gene cassette and a second component supplying gvpA together with constitutive rtTA. The H−R+ line used in the phantom experiments and tracked in **Fig. 6** carried pSS-16 (pLenti-TRE3G-AnaN –V–EmGFP), comprising P2A-separated gvpN, gvpJ, gvpK, gvpF, gvpG, gvpW and gvpV followed by EmGFP and lacking WPRE, together with mL-82 (pLenti-TRE-A-WPRE-mEF1a-rtTA), which comprised TRE3G-driven gvpA–WPRE and constitutive minimal-EF1α-driven rtTA. The H+R+ line used in **Fig. 5** carried mL-88 (pLenti-TRE3G-AnaN-V-EmGFP-WPRE-mEF1a-BSD), comprising the same TRE3G-driven, P2A-separated gvpN–J–K–F–G–W–V–EmGFP cassette followed by WPRE and a minimal-EF1α–BSD selection cassette, together with mL-82 and mL-77 (pLenti-EF1a-hHER2-RPBSA-PuroR). mL-77 comprised full-length EF1α-driven hHER2 and an RPBSA–PuroR selection cassette. The H+R− reporter-negative background line used in the reciprocal **Fig. 6** model carried mL-77 without the GV reporter vectors. Accordingly, H+ and H− denote the presence or absence, respectively, of the ectopic hHER2 vector, whereas R+ and R− denote the presence or absence of the complete inducible GV reporter system; these labels do not imply complete absence of endogenous HER2.^14,18^

### Cell line generation

All lentiviral functional titers were measured in HEK-293T cells. For stable MDA-MB-231 line generation, target cells were plated in six-well plates and allowed to adhere for 24 h before vector exposure in complete medium containing 10 µg ml⁻¹ polybrene. Virus-containing medium was removed after transduction, and cells were transferred to expansion culture within 24–36 h and maintained in surface-treated T-75 flasks. H−R+ cells were co-transduced with pSS-16 and mL-82 using functionally titrated stocks. H+R− cells were transduced with mL-77 at a multiplicity of infection (MOI) of 4 and selected with puromycin. H+R+ cells were generated from the puromycin-selected H+R− population by transduction with mL-88 at MOI 1 and mL-82 at MOI 4.^18^

Antibiotic selection was matched to the resistance cassette. MDA-MB-231 cells were cultured with 2 µg ml⁻¹ puromycin, 200 µg ml⁻¹ hygromycin or 10 µg ml⁻¹ blasticidin, as applicable. Corresponding selection concentrations for HEK-293T cells were 3 µg ml⁻¹ puromycin, 300 µg ml⁻¹ hygromycin or 10 µg ml⁻¹ blasticidin. Cells were cultured and serially passaged under selection for at least 2 weeks.

The GV-expressing HEK-293T line was generated by lentiviral rtTA delivery followed by PiggyBac integration of the reporter cassettes. Cells were first transduced with mL-47 (pLenti-EF1a-rtTA-P2A-HygR-WPRE) at MOI 0.2 to favor single-copy integration of the rtTA cassette and selected with hygromycin. The resulting HEK-mL47 cells were transfected with mT-45, the PiggyBac counterpart of the mL-88 reporter cassette, and mT-61 (pPB-TRE3G-A-WPRE-bGH) at a 1:4 molar ratio, together with PiggyBac transposase at a 2.5:1 mass ratio of transgene DNA to transposase DNA. Cells were then maintained under hygromycin and blasticidin selection.^18^

After selection, reporter-positive MDA-MB-231 and HEK-293T populations were induced with 1 µg ml⁻¹ doxycycline for 48 h before fluorescence-activated cell sorting. Parental cells of the corresponding lineage established the GFP and side-scatter gates, and GFP-positive cells with side scatter above the parental distribution were collected. Sorted cells were expanded in tetracycline-free medium. For the MDA-MB-231 populations, HER2 expression was verified by flow cytometry and single-cell confocal fluorescence imaging before implantation. This induction interval was used only during cell-line establishment; experimental induction before phantom imaging is described below. Within each imaging experiment, induced and uninduced samples were derived from the same stable population and underwent identical preparation and acquisition.^14,18^

### Doxycycline induction for in vitro experiments

Reporter expression was induced by adding doxycycline to culture medium at 1 µg ml⁻¹. Medium containing fresh doxycycline was replaced every 24 h, and cells were harvested after six days of induction for the phantom experiments. Uninduced cells from the same reporter-carrying preparations were cultured in parallel without doxycycline and served as reporter-negative controls. The dose, daily replacement and induction interval were selected from prior characterization of MDA-MB-231 GV expression.^4,14^

### Agarose phantom preparation

Multiwell phantoms were cast from 1% (w/v) agarose in phosphate-buffered saline (PBS). On imaging day, induced and uninduced cells were detached with Accutase for 15 min and released with complete medium, passed through a 40-µm cell strainer to remove aggregates, counted and pelleted at 300g for 6 min. Reporter and background cells were resuspended separately at twice their final loading concentrations. Each 40-µl well contained 20 µl of 2% low-melting-point agarose in PBS at 42 °C, 10 µl of reporter-cell suspension and 10 µl of background-cell suspension, yielding 1% agarose and the cell numbers specified below. Samples were mixed immediately before loading, transferred without bubbles, allowed to solidify and submerged in PBS for imaging.^4,14^

### Cell composition of phantom wells

HEK-293T wells contained 600 induced reporter-carrying cells mixed with 12,000 uninduced reporter-negative cells. MDA-MB-231 wells contained 4,500 induced reporter-carrying cells mixed with 13,500 uninduced reporter-negative cells. Five induced and five uninduced control wells were prepared for each cell line. Cell chirp and cross-propagating wave amplitude modulation (xAM) acquisitions were interleaved in the same wells, at the same imaging planes, to permit paired comparisons. All 20 wells were retained in the complete paired dataset.

### Optical estimation of GV-positive cells

Cells were imaged on an Axio Observer A1 inverted microscope (Zeiss) with an A-Plan 10×/0.25 Ph1 objective and an AxioCam ICm1 monochrome camera (1,388 × 1,038 pixels, 14-bit; 1 × 1 binning) controlled by ZEN core v.2.5.74.0. For each field, Hoechst and GFP were acquired as separate single-channel epifluorescence images, and a transmitted-light phase-contrast image was acquired with HAL 100 halogen illumination. Fields were sampled across individual wells. Nuclei were segmented from the Hoechst channel with StarDist, and each segmented cell was classified for GFP fluorescence and, from the corresponding phase-contrast image, for the dark refractile intracellular inclusions characteristic of gas-vesicle (GV) expression. Non-adherent dead cells were excluded. The fraction of cells containing optically visible GVs was multiplied by the number of reporter-carrying cells loaded into the corresponding well to obtain the optical GV-positive estimate used for comparison with acoustic punctum counts. GFP-positive estimates were calculated analogously. Optical and acoustic counts were not treated as one-to-one registered measurements.

### Ultrasound instrumentation and acquisition geometry

Ultrasound data were acquired with a Verasonics Vantage NXT scanner and a GE 6-24D linear-array probe at a 50-MHz RF sampling frequency. The probe comprises 192 elements with a 0.135-mm pitch. To reduce artifacts from outside the imaging field, edge elements were disabled for both transmit and receive. For phantom imaging, 30 elements on each side were disabled, leaving 132 active elements. The 65-element transmit subaperture was stepped laterally by one element at a time across the active aperture, yielding 68 raylines. For in vivo imaging, 20 elements on each side were disabled, leaving 152 active elements; the same stepping scheme yielded 88 raylines. For both phantom and mouse imaging, the probe was secured in a custom 3D-printed holder mounted on a motorized three-axis positioner (Genmitsu 3020-PRO Ultra, SainSmart). B-mode, cell chirp and xAM were acquired without moving the sample. Volumes were acquired at 100-µm elevational steps in phantoms and 200-µm steps in mice; the coarser in vivo spacing shortened volumetric acquisition and anesthesia time. At each plane, one B-mode image and four cell chirp and four xAM images were acquired, with cell chirp and xAM interleaved. Each cell chirp and xAM image was formed on the hardware by averaging 25 repeated transmit–receive acquisitions at the same receive-buffer location, following the averaging approach described previously.^14^ The four images within each pulse sequence were not averaged together: the third cell chirp image and the third xAM image were used consistently for all primary analyses, whereas all four images were analyzed separately to assess repeatability. Reconstructed sampling intervals were 15.9 µm axially and 100 µm laterally. Phantoms were submerged in PBS, and anesthetized mice were coupled to the transducer with ultrasound gel.

### Cell chirp and xAM pulse sequences

Cell chirp used the same cross-propagating-wave amplitude-modulation framework as xAM; both sequences transmitted at 20° through a 65-element subaperture. xAM acquisitions used a 15.625-MHz transmit frequency. Because the default scanner-generated chirp produced unequal cycle amplitudes, a duty-cycle-loaded chirp was implemented as a custom tri-level waveform using Verasonics CStates on the scanner’s 500-MHz transmit clock. Each cell chirp event comprised one full-amplitude and two half-amplitude transmissions of the same 13-cycle downward sweep. The design frequency decreased linearly by cycle from 10.5 to 7.0 MHz. Each cycle consisted of an undriven interval, a positive-drive interval, a second undriven interval and a negative-drive interval, with equal positive- and negative-drive durations. Because the drive voltage was fixed within each transmit event, the amplitude of each cycle was controlled through its duty cycle, defined as the fraction of the period spent driven. Per-cycle duty cycles were manually assigned using duty-cycle modeling to compensate for the frequency dependence of transmitted pressure and produce an approximately uniform peak-positive pressure across the sweep. From the first to the last cycle, the programmed duty cycles were 70%, 70%, 62%, 50%, 50%, 55%, 60%, 65%, 70%, 90%, 90%, 89% and 89%, with the longest drive intervals assigned to the lowest frequencies. Each interval was rounded to a whole number of 2-ns clock periods, yielding realized cycle frequencies from 10.42 to 6.58 MHz and realized duty cycles from 50.0% to 90.9%. The same duty-cycle-loaded waveform was used for phantom and in vivo imaging and was characterized by fiber-optic hydrophone as described below. Full- and half-amplitude echoes were combined to suppress linear scattering while retaining nonlinear GV responses. xAM and B-mode were acquired as paired comparators. Echoes were received across the transducer bandwidth, including the fundamental and second-harmonic bands. Cell chirp and xAM were acquired at peak-positive pressures of 0.393 MPa and 0.513 MPa, respectively.

### Imaging pressures and GV-collapse controls

Cell chirp and xAM imaging was performed at fixed peak-positive pressures of 0.393 MPa and 0.513 MPa, respectively. These pressures were below the empirically observed onset of irreversible GV signal loss and were used for both phantom and in vivo imaging. The two sequences were not voltage- or pressure-matched because their durations and spectra differed. After the final induced tumor scan, a separate focused suprathreshold sequence was applied deliberately to collapse GVs, and the same tumor volume was reacquired within minutes as a loss-of-signal control. Routine longitudinal acquisitions did not use collapse as the detection event.

### Fiber-optic hydrophone data acquisition and waveform analysis

Transmit pressures and source waveforms were measured in a water tank with a calibrated Precision Acoustics Fibre-Optic Hydrophone System (serial PAFOH44) using sensing fibre TFS-5899-01. The fibre tip was positioned at 5-mm depth and aligned to the acoustic maximum. The same GE 6-24D probe, transmit aperture and programmed cell chirp and xAM sequences used for imaging were used for hydrophone measurements. Source-waveform shape was recorded at a matched 3.0-V drive, and the fibre-optic-hydrophone calibration was used separately to set the sequence-specific imaging peak-positive pressures; no needle hydrophone was used for the reported measurements. For display, cell chirp and xAM traces were normalized independently to their maximum absolute amplitude. Measured durations were 1.76 and 0.49 µs, respectively, and the corresponding −6-dB frequency bands were 7.3–9.8 and 8.2–16.3 MHz. For the waveform-shape comparison, each single full-amplitude pressure trace was normalized to the same peak absolute pressure and integrated as ∫p(t)²dt. The resulting cell chirp- to-xAM squared-pressure time-integral ratio was 9.6; this quantity is not the energy ratio of the complete amplitude-modulation acquisitions at their unequal imaging pressures. The transmitted cell chirp spectrum was also inspected in the second-harmonic receive band to verify that harmonic-band sample responses were not explained by appreciable harmonic content in the transmitted waveform.^14^

### In vitro imaging of engineered MDA-MB-231 and HEK-293T cells

Solidified cell-containing phantoms were submerged in PBS and centered at the calibrated imaging depth. Each induced well and its uninduced control were imaged with cell chirp, xAM and B-mode using the fixed acquisition settings described above. The same planes were used for both nonlinear sequences. The MDA-MB-231 preparation linked the in vitro assay to the orthotopic tumor studies; the HEK-293T preparation provided the low-cell-number benchmark. Sequence comparisons used well-level paired measurements, and repeated-readout experiments retained the same well and imaging position over successive acquisitions.^4,14^

### Beamforming, harmonic filtering and matched filtering

Amplitude-modulated channel data were beamformed to form full-band and harmonic-band reconstructions. The harmonic reconstruction was filtered and matched to an eight-cycle frequency-encoded template spanning 12.5–23 MHz. Matched filtering compressed the encoded nonlinear response into a localized peak. At each axial position, local reference levels were calculated from the full-band and harmonic-band envelopes in windows extending ±22 and ±11 samples, respectively, with the central response excluded. Let E denote the matched-filter envelope and B the geometric mean of these two local reference levels. The detector statistic was S = 20 log₁_0_(E) + 5[20 log₁_0_(E/B)]. S is a composite decibel-scale score used for detection and is not a calibrated acoustic-pressure or echo-amplitude measurement.

### Reporter detection and fixed thresholds

One detector configuration was used for all phantom datasets and one configuration for all in vivo datasets. The cell chirp threshold was 79.6 dB on the composite detector-score scale for phantom data and 74.8 dB on the same scale for in vivo data. Each cell chirp threshold was the lowest value at which pooled reporter-negative controls gave no more than a fixed number of false detections, and was then held fixed across the corresponding dataset. For phantom data, the controls were the five uninduced wells of each cell line, and the limit was the number of detections that the earlier version of the analysis pipeline gave in the same wells (74 for HEK-293T and 44 for MDA-MB-231). The MDA-MB-231 limit gave the higher threshold, 79.6 dB, which was used for both cell lines; at this threshold the control wells gave 57 (HEK-293T) and 34 (MDA-MB-231) detections before de-duplication. For in vivo data, the controls were the four post-collapse tumor volumes of the induction cohort, and the limit was 72 detections, carried over from an earlier version of the detector; 74.8 dB gave 68 detections when it was set. The threshold was not re-derived after the counting regions and analyzed imaging planes were finalized; in the final counts, the four post-collapse volumes gave 37 detections at 74.8 dB. xAM detections used the previously described analysis pipeline^14^ with a fixed 40-dB envelope threshold, which was not re-derived for this study. Because the two thresholds were set differently, in vivo detection was also compared at a matched false-detection rate (**Extended Data Fig. 5n,o**). For both sequences, a detection required at least three contiguous axial samples above threshold. Threshold sweeps used for descriptive performance comparisons were analyzed separately from the fixed thresholds used for the principal counts and were not treated as held-out validation.

### Detection localization, counting and three-dimensional rendering

The position of each threshold-positive cell chirp response was refined with a three-point parabolic fit to the detector-statistic maximum. Fitting was performed only after threshold crossing and therefore affected peak localization and display, not whether a detection was counted. In the phantom data, detections in adjacent elevation planes (100 µm apart) were linked only between neighboring planes when they lay within one lateral column and either their axial centers were within three samples or their axial extents overlapped by at least 25%. Each detection was assigned at most one partner in the next plane, with candidate matches accepted greedily in order of lateral distance and then axial distance. Links were chained across consecutive planes, and the strongest detection in each chain was retained. No within-plane merging was performed because the three-sample axial-run rule already yielded one detection per run. This procedure removed 51% of detections in induced HEK-293T wells, 45% in induced MDA-MB-231 wells and 48% across the two induced preparations, compared with 0–4% in uninduced wells; 55 of 57 HEK-293T and 34 of 34 MDA-MB-231 control detections were retained. In vivo volumes were acquired at 200-µm elevational steps, and detections were counted separately in each plane and summed without cross-plane consolidation. A source spanning adjacent planes could therefore contribute more than one detection. A manually outlined tumor-containing counting region was applied to each volume, and the identical region was used for paired cell chirp and xAM data and for pre- and post-collapse comparisons. Counts were obtained from detection coordinates and were not normalized by counting-region volume. Three-dimensional views were generated in napari by overlaying fitted puncta on the B-mode volume.

### Animals and tumor-cell implantation

All animal procedures were approved by the California Institute of Technology Institutional Animal Care and Use Committee under protocol no. 1721. Female NSG (NOD.Cg-Prkdcscid Il2rgtm1Wjl/SzJ) mice aged 8–16 weeks were obtained from The Jackson Laboratory (stock no. 005557) and used to establish orthotopic human breast-cancer xenografts. MDA-MB-231 populations were washed once with warm PBS, incubated with Accutase for 15 min and released with complete medium. Cells were passed through a 40-µm cell strainer to remove aggregates, counted with trypan-blue exclusion, centrifuged at 500g for 5 min and washed twice in PBS. The reporter and background populations were resuspended separately at 1 × 10⁶ cells ml⁻¹, combined at the required cell numbers, pelleted again at 500g for 5 min and resuspended in 300 µl of ice-cold Matrigel diluted 1:1 with PBS, providing a 300-µl injection volume per tumor. Cell suspensions were remixed immediately before loading into 1-ml insulin syringes; loaded syringes were kept needle-down on ice, and each suspension was injected slowly and unilaterally into the left lower mammary fat pad to minimize bubble formation. Mice were induced with 5% isoflurane and maintained at 3.5% isoflurane through a nose cone during implantation. The induction cohort received 2 × 10⁴ reporter-positive cells together with 1 × 10⁶ reporter-negative cells. A separate reporter-input experiment used either 2 × 10⁴ or 4 × 10^3^ reporter cells together with 1 × 10⁶ reporter-negative cells (n = 5 tumors per group). For ultrasound imaging, mice were induced with 5% isoflurane and maintained at 1.5–2% isoflurane through a nose cone. Tumor dimensions were measured with calipers, and volume was calculated as π/6 × length × width × height.

### Doxycycline induction and longitudinal in vivo imaging

For in vivo reporter induction, doxycycline was prepared in sterile saline at 1 mg ml⁻¹, and each mouse received a fixed 150-µl intraperitoneal dose containing 150 µg doxycycline. The fixed dose matched previous GV-reporter tumor work, but the present study used once-daily dosing. ^4^ In the daily induction experiment, four tumors were imaged before doxycycline and again 24 h after each of three consecutive daily doses. The same tumors were followed at each time point. After the third induced scan, GVs were deliberately collapsed and the tumors were re-imaged within minutes. The reporter-input cohorts were imaged on day 7.

### Human T-cell isolation, activation and expansion

Human T cells were isolated from cryopreserved peripheral blood mononuclear cells from a single donor (STEMCELL Technologies); donor material was not pooled. Vials were thawed into prewarmed RPMI-1640 containing 0.1 mg ml⁻¹ DNase I, diluted dropwise to 25 ml and centrifuged at 400g for 10 min at 25 °C. Cells were passed through a 40-µm strainer, counted with trypan-blue exclusion and enriched by negative selection with the EasySep Human T Cell Isolation Kit (STEMCELL Technologies) at approximately 5 × 10⁷ cells ml⁻¹. Complete T-cell medium was prepared from 500 ml RPMI-1640, 50 ml fetal bovine serum, 5 ml GlutaMAX and 5 ml penicillin–streptomycin. Recombinant human IL-2 IS (Miltenyi Biotec, cat. no. 130-097-743) was reconstituted by adding 500 µl molecular-grade water to a 50-µg vial, yielding a 100 ng µl⁻¹ stock; 5- or 10-µl aliquots were stored at −30 °C. Cells were activated with washed Dynabeads Human T-Activator CD3/CD28 beads (Gibco; 25 µl bead suspension per milliliter of culture). Cultures were maintained in 45 ml complete medium per 600-ml suspension-culture flask at 0.3–1 × 10⁶ cells ml⁻¹ and supplemented with IL-2 at 10 ng ml⁻¹ (4.5 µl stock per 45-ml culture). For cell counting and medium renewal, cultures were mixed gently, a 10-µl aliquot was combined 1:1 with trypan blue, and the remaining cells were centrifuged at 350g for 5 min at room temperature before resuspension in fresh IL-2-supplemented medium. Activation beads were removed magnetically on day 5, and cells were expanded for 10 days. Before injection, cells were counted for viability, washed and resuspended in sterile PBS.^15^

### HER2 × CD3 bispecific-antibody administration and longitudinal treatment imaging

For the directly targeted model, tumors contained 2 × 10⁴ reporter-positive cells carrying ectopic hHER2 among 1 × 10⁶ reporter-negative cells without ectopic hHER2 (n = 5 mice per group). For the antigen-negative model, tumors contained 2 × 10⁴ reporter-positive cells without ectopic hHER2 among 1 × 10⁶ reporter-negative cells carrying ectopic hHER2 (n = 5 treated and n = 4 control mice). Immediately before administration, expanded human T cells were passed through a 40-µm cell strainer to remove aggregates, pelleted, washed twice in sterile PBS and resuspended at 1 × 10⁷ cells per 150 µl. After the second weekly imaging session, each mouse received 1 × 10⁷ T cells in 150 µl sterile PBS by tail-vein injection. Runimotamab, a HER2 × CD3 T-cell-dependent bispecific antibody (TDB), was diluted in sterile saline to a working concentration of 100 µg ml⁻¹, and 100 µl (10 µg) was administered intraperitoneally per mouse. The first dose was given 30 min before T-cell transfer, and dosing continued every 48 h during the treatment window.^11,15^ Control mice received T cells without TDB. Tumors were imaged at two pretreatment and two post-treatment weekly time points using the same cell chirp, xAM and B-mode acquisition workflow.

### Tumor harvest, fixation and immunofluorescence staining

Tumors were collected immediately after the final ultrasound session and fixed overnight in 10% formalin at 4 °C with continuous rotation. Tissue was cryoprotected sequentially in 15% sucrose for 24 h and 30% sucrose for 48 h, embedded in O.C.T. compound, stored at −80 °C and cryosectioned at 25-µm thickness. Sections were washed three times for 5 min in PBS with gentle agitation. Sections were permeabilized with 0.1% Triton X-100 in PBS for 10 min at room temperature, washed three times for 5 min in PBS and blocked with 5% bovine serum albumin in PBS for 60 min at room temperature. After blocking, sections were washed three times for 5 min in PBS. Alexa Fluor 647-conjugated anti-human CD8 antibody (BioLegend, clone SK1, cat. no. 344726) was diluted in 1% bovine serum albumin in PBS and incubated overnight at 4 °C in a humidified chamber. Sections were washed three times for 5 min in PBS, mounted in DAPI-containing antifade medium, coverslipped and cured overnight at room temperature in the dark. GFP was detected from the endogenous co-reporter without antibody amplification.^15^

### Confocal microscopy and image quantification

Whole-section mosaics were acquired as single optical planes on a Leica STELLARIS 8 FALCON confocal microscope using an HC PL APO CS2 20×/0.75 IMM objective with distilled-water immersion. Fluorescence channels were acquired sequentially at 12-bit depth with a 2.00-Airy-unit pinhole (at 580 nm). GFP and CD8 acquisition settings were held constant across datasets included in the respective quantitative analyses. One section per animal was analyzed within DAPI-defined tumor boundaries, excluding a 150-µm peripheral margin. Necrotic regions larger than 0.02 mm² and a 60-µm margin surrounding each region were excluded. GFP-positive cells were identified with a fixed intensity threshold. CD8-positive objects were detected with a fixed classifier and retained when their projected area was 25–100 µm².^15,16^

### Whole-tumor clearing and light-sheet microscopy

For the whole-tumor comparison in **Fig. 4j**, an excised tumor was fixed in 10% neutral buffered formalin for 12 h, cryoprotected in 15% sucrose for 24 h and 30% sucrose for 48 h, embedded in O.C.T. and stored at −80 °C. Excess O.C.T. was trimmed while cold, and the intact specimen was washed repeatedly in cold PBS, including overnight at 4 °C. A modified SHIELD workflow^26^ was used to preserve endogenous GFP: 48 h at ≤4 °C in SHIELD-OFF solution (50% SHIELD Epoxy, 25% SHIELD Buffer and 25% water), followed by 24 h in SHIELD-ON solution at 37 °C. The tumor was passively delipidated with gentle agitation in approximately 5 ml of LifeCanvas Delipidation Buffer at 45 °C for 4 d, with a buffer change after 48 h, and washed in PBS containing 1% Triton X-100.

Organic-solvent delipidation followed a modified FDISCO procedure^27^. PBSQ contained 25% (v/v) Quadrol (Sigma-Aldrich, 122262) in PBS. The tumor was incubated in 50%, 70%, 80%, 95% and fresh 95% tetrahydrofuran (THF) in PBSQ for 1 h each, then in 100% dichloromethane for 30 min and fresh 100% dichloromethane for approximately 15 h. The THF series was reversed for rehydration, using the same incubation times. All THF and dichloromethane steps were performed at 4 °C in the dark. The tumor was washed for at least 24 h in PBS containing 0.02% sodium azide with multiple buffer exchanges, and clearing was completed by refractive-index matching in EasyIndex (LifeCanvas Technologies).

For imaging, the tumor was immersed in an ethyl cinnamate/mineral oil mixture adjusted to a refractive index of approximately 1.52. Endogenous GFP was imaged without antibody amplification on a LaVision BioTec UltraMicroscope II controlled by ImSpector Pro 7.6.4, using a 2× objective at 0.63× zoom (1.26× total magnification), 488-nm excitation and a 525/50-nm emission filter. Bilateral light-sheet illumination was used, with the two sides merged in blend mode. An Andor camera recorded 2,293 planes spanning 4.58 mm at 16-bit depth, with 30-ms exposures and a voxel size of 5.159 × 5.159 × 2.0 µm. Five optical planes were reoriented for display to align gross tumor anatomy with the ultrasound rendering; GFP puncta were displayed at least 2.5-fold above local background. The modalities were not pointwise registered.

### Statistical analysis

Wells were the independent units for phantom experiments and animals were the independent units for in vivo experiments; puncta, imaging planes and microscopy fields were nested measurements and were not treated as independent replicates. All tests were two-sided with α = 0.05. Detection counts were analyzed after log_10_ transformation, and fold differences for these analyses are ratios of geometric means. Phantom cell chirp and xAM detection yields were compared within wells using paired t-tests on log-transformed counts. Well-level paired summaries of axial extent, signal-to-background ratio and co-detected versus cell chirp-only puncta were compared with Wilcoxon signed-rank tests. For the induction series, the first-dose response was the within-animal ratio of the first induced count to baseline, and the daily factor was the square root of the ratio of the third to the first induced count; one-sample t-tests were applied to the corresponding log ratios. The day-7 reporter-input comparison and weekly treatment-group comparisons used Welch’s t-tests on log-transformed counts with animals as replicates. Endpoint tumor-volume and microscopy comparisons used two-sided Welch’s t-tests on the per-animal measurements specified in the figure legends. Associations between acoustic counts and endpoint GFP-positive density were evaluated by linear regression of log-transformed values, with tumor model and, where indicated, treatment group included as covariates. P values were not adjusted for multiple comparisons. Significance symbols denote *P < 0.05, **P < 0.01, ***P < 0.001 and ****P < 0.0001. Exact sample sizes and P values are reported in the figure legends. Analyses were performed in GraphPad Prism 10 and Python.

### Data and code availability

Source data, ultrasound acquisition sequences, reconstruction code and detector code will be deposited in public repositories before publication. Additional raw data will be available from the corresponding author upon reasonable request.

## Author contributions

R.N. and M.G.S. conceived the study. Z.J., M.D., and R.N. generated and sorted the cell lines. T.F. assisted R.N. with tissue culture and preparation of cryopreserved cell stocks. R.N. performed tissue culture, the in vitro and in vivo experiments, histology and microscopy, and H.A. performed the intravenous T-cell injections. K.L. provided training in T-cell harvesting and input on the study design. R.N. processed and analyzed the data. R.N. and M.G.S. wrote the manuscript with input from all other authors. M.G.S. supervised the research.

## Acknowledgements

We thank Mickael Tanter and Grace Shin for their contributions to this work; Shirin Shivaei, Hao Shen, George Daghlian and Daniel Tang for helpful discussions; and Zhongying Wang of the Caltech Biological Imaging Facility for microscopy support. This research was supported by the National Institutes of Health (R01-EB018975 to M.G.S.) and the Chan Zuckerberg Initiative. M.G.S. is an Investigator of the Howard Hughes Medical Institute. **Figs. 1, 3a, 4a, 5a** and 6a were created in BioRender (BioRender.com/idsurqg; BioRender.com/x9j1mpy).

## Extended Data

**Extended Data Fig. 1.**
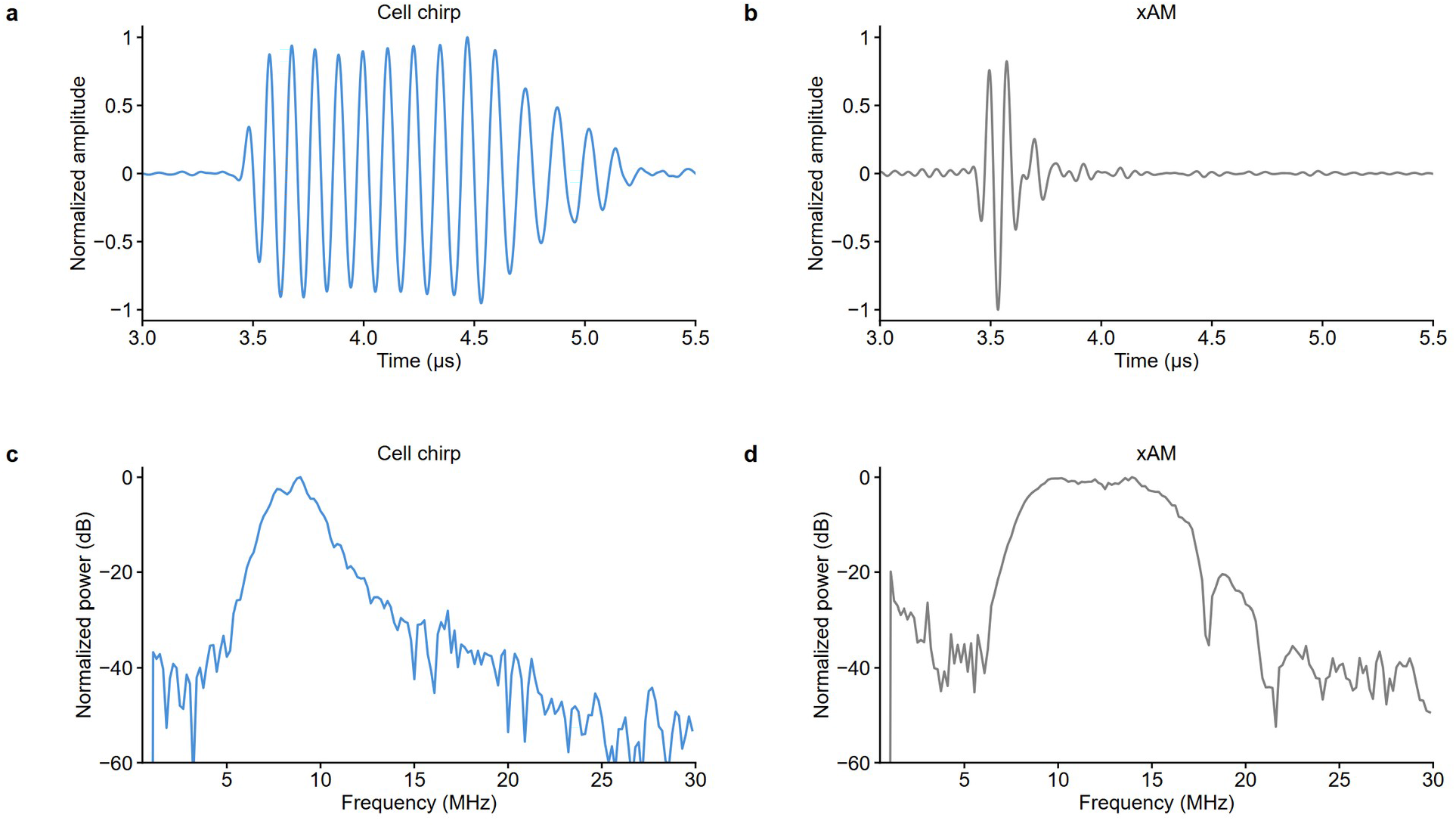
Cell chirp extends excitation while maintaining low harmonic-band transmission. a,b, Hydrophone-measured cell chirp (a) and xAM (b) transmit waveforms at 5-mm depth using the same linear-array transducer and aperture used for imaging. Each waveform was normalized to its own maximum absolute amplitude and plotted on common axes. Cell chirp extended the measured transmission to 1.76 µs, compared with 0.49 µs for xAM. c,d, Power spectra of the cell chirp (c) and xAM (d), normalized to the maximum power of the corresponding waveform. The measured −6-dB bands were 7.3–9.8 MHz for cell chirp and 8.2–16.3 MHz for xAM. The cell chirp spectrum fell by approximately 35–40 dB in the second-harmonic receive band, supporting attribution of harmonic-band reporter echoes to the sample rather than appreciable nonlinear content in the transmitted waveform.

**Extended Data Fig. 2.**
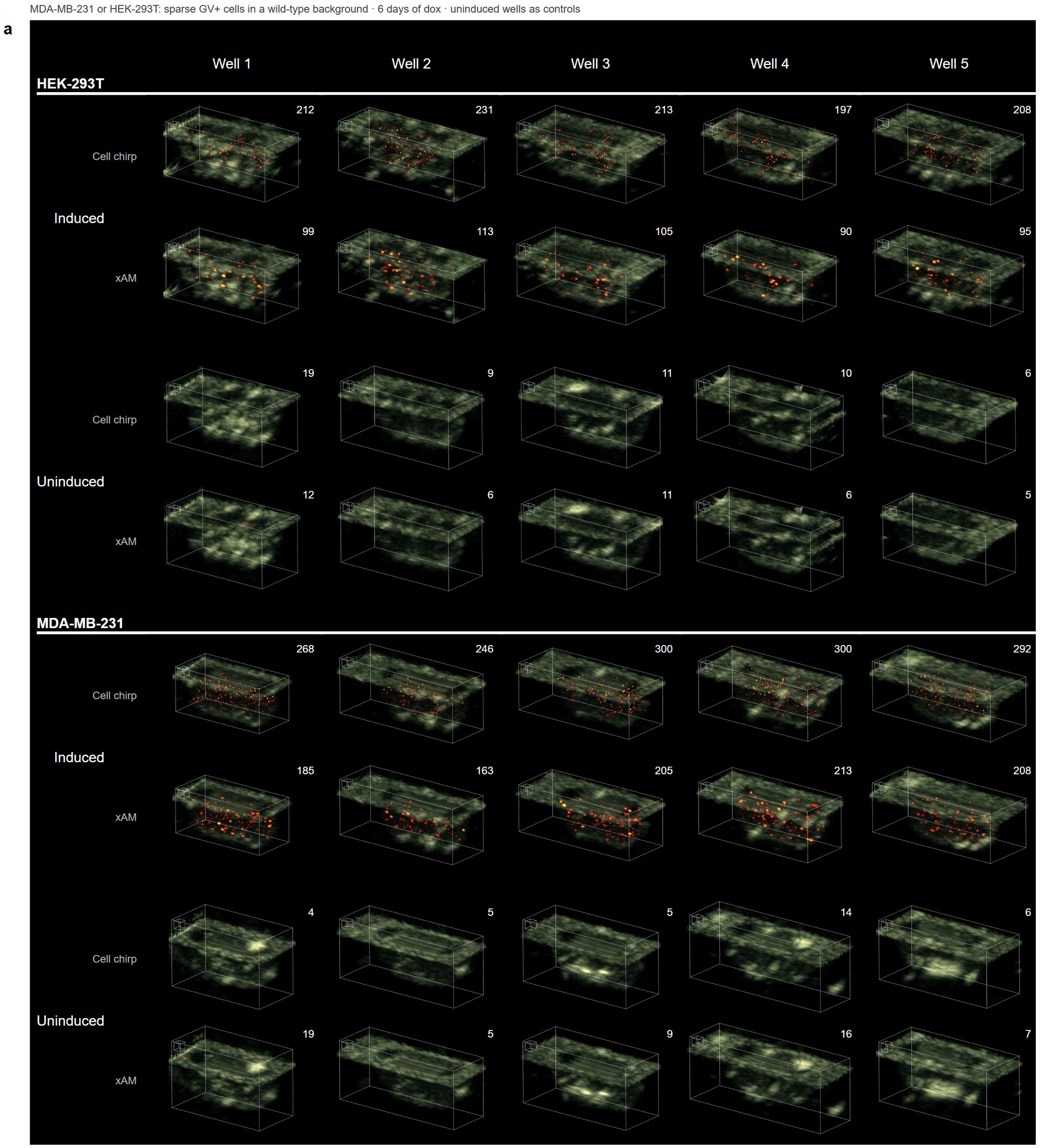
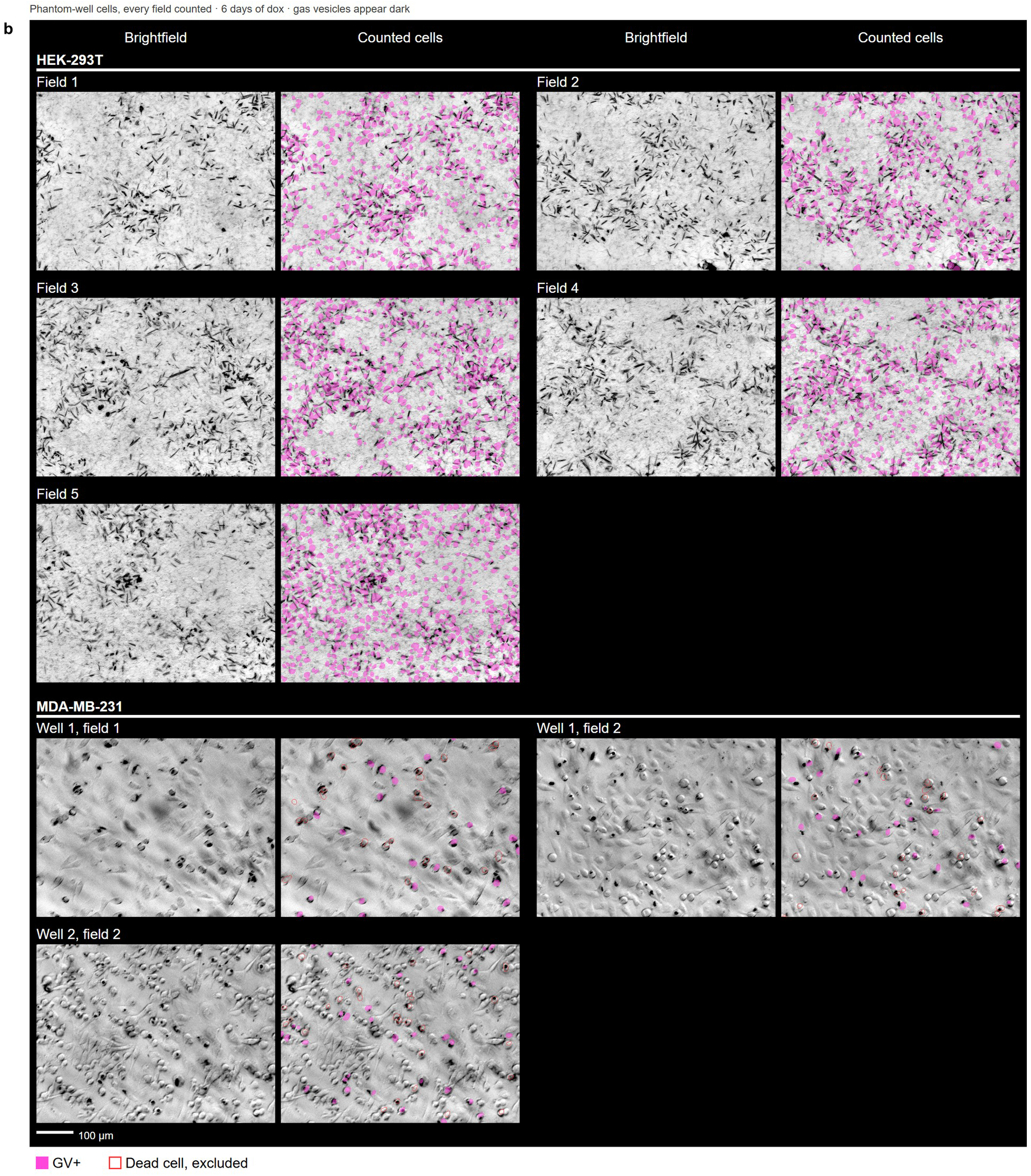
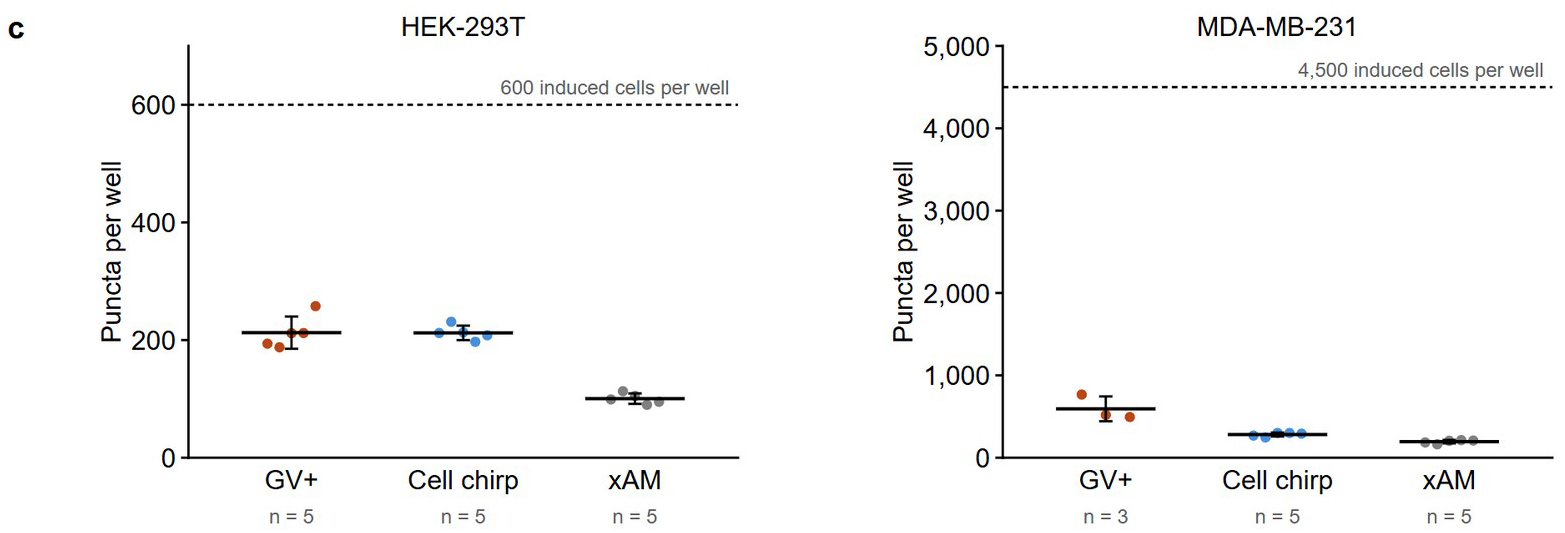
Complete phantom datasets and optical benchmarks of reporter recovery. a, Complete paired phantom dataset. Volumetric cell chirp and xAM detections are shown for five induced and five uninduced control wells from each of the MDA-MB-231 and HEK-293T preparations (20 wells; 40 volumes). Each column is one well, with cell chirp and xAM in adjacent rows. Induced wells were imaged after six days of doxycycline treatment; uninduced wells from the same preparations served as reporter-negative controls. All renderings use the same physical scale, crop and B-mode display window; cubes indicate 0.5 mm. Numbers are absolute detection counts for each well. Cell chirp produced more induced-well detections in all five wells from both cell lines while retaining low counts in uninduced controls. The representative volumes in Fig. 3b are MDA-MB-231 induced well 1 and control well 2, and HEK-293T induced well 2 and control well 2. b, Brightfield fields used to estimate the number of reporter cells containing optically visible GVs after six days of doxycycline induction. Each unannotated image is paired with its counted image. Magenta denotes GV-positive cells; red denotes non-adherent dead MDA-MB-231 cells excluded from counting. Five HEK-293T fields and three MDA-MB-231 fields are shown. Scale bar, 100 µm. c, Optical GV-positive estimates and acoustic detections per induced phantom well. Points denote individual counted fields or acoustic wells; bars show mean ± s.d.; dashed lines mark the number of induced cells added per well. GFP-positive estimates were 362.0 ± 20.0 cells per HEK-293T well (60.3% of the 600 induced cells) and 4,367.5 ± 117.0 cells per MDA-MB-231 well (97.1% of the 4,500 induced cells). Optically visible GV-positive estimates were 212.6 ± 27.4 and 593.4 ± 150.6 cells per well for HEK-293T and MDA-MB-231 preparations, respectively. Cell chirp detected 212.2 ± 12.3 and 281.2 ± 23.6 puncta per well, whereas xAM detected 100.4 ± 8.9 and 194.8 ± 20.7 puncta per well. Optical estimates used five HEK-293T fields and three MDA-MB-231 fields; acoustic measurements used five wells per cell line.

**Extended Data Fig. 3.**
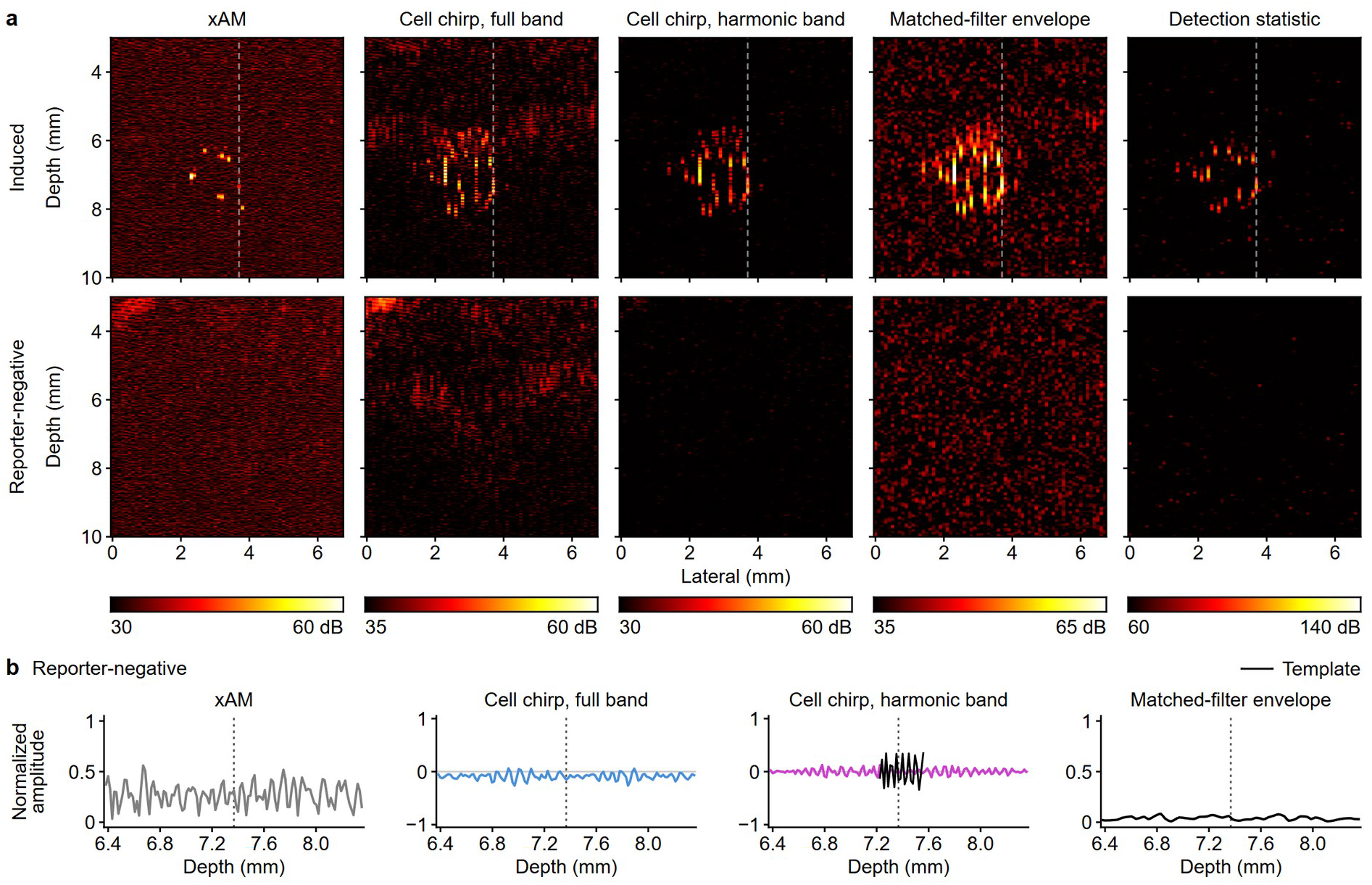
Induced and reporter-negative controls verify reporter-specific signal recovery in phantoms. a,b, Signal-recovery controls in induced and reporter-negative MDA-MB-231 phantoms. a, Five processing stages for an induced preparation (top) and a reporter-negative control (bottom), shown at matched color limits: xAM, cell chirp full band, cell chirp harmonic band, matched-filter envelope and detection statistic. The dashed line marks the induced rayline analyzed in Fig. 3g; b shows the reporter-negative control at the same rayline. Detections required at least three contiguous axial samples above the fixed 79.6-dB in vitro threshold. b, The reporter-negative response remained below threshold under the induced-trace normalization and template placement.

**Extended Data Fig. 4.**
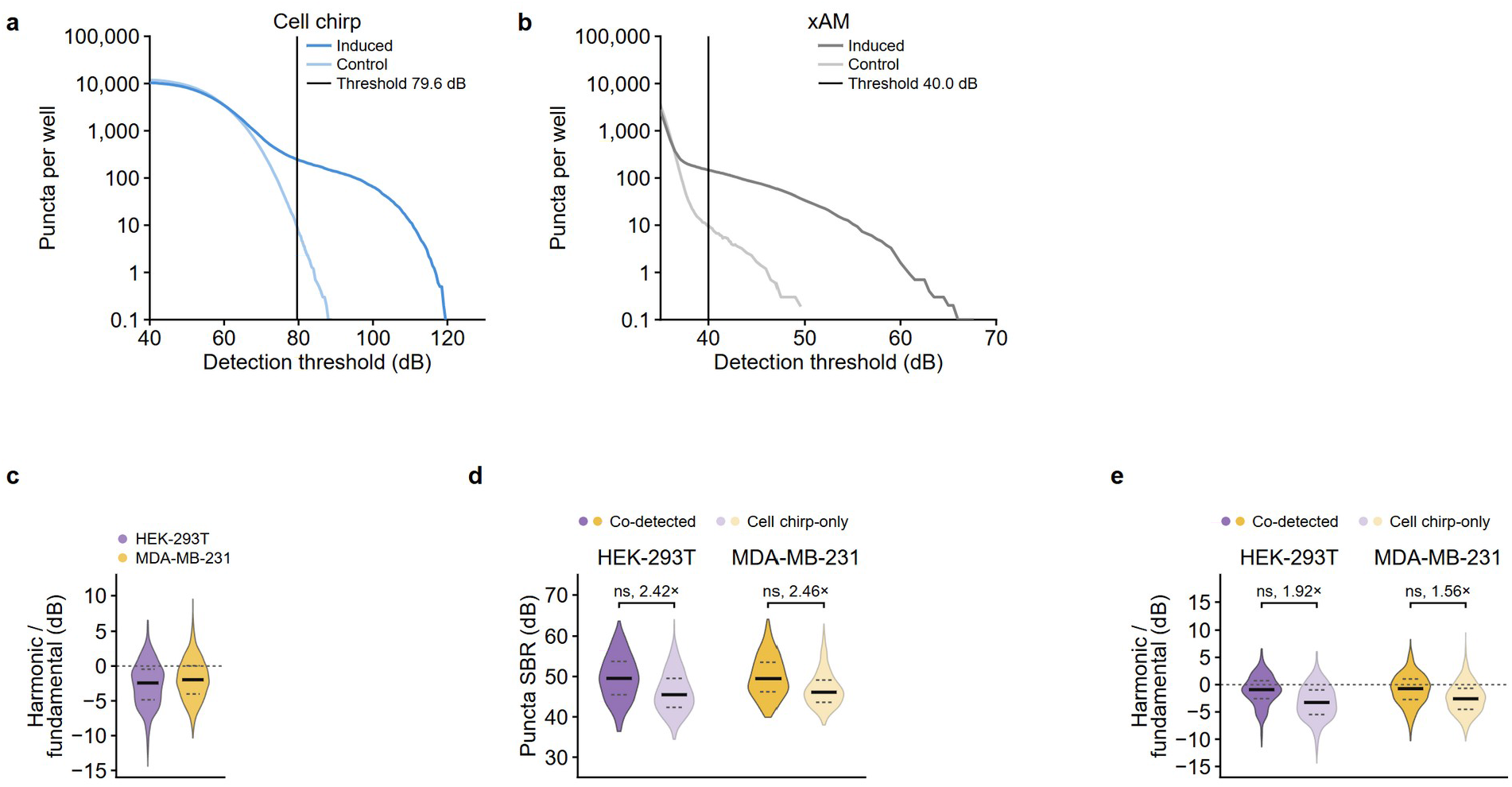
In vitro threshold selection and weak-signal phenotypes. a,b, Phantom threshold sweeps for cell chirp (a) and xAM (b). Curves show mean de-duplicated puncta per well in ten induced wells and ten uninduced control wells pooled across HEK-293T and MDA-MB-231 preparations. Vertical lines mark the fixed detection thresholds (79.6 dB for cell chirp and 40.0 dB for xAM). c, Harmonic-to-fundamental mean power ratio for cell chirp puncta (n = 1,061 HEK-293T and 1,406 MDA-MB-231 puncta). d,e, Signal-to-background ratio (d) and harmonic-to-fundamental power (e) for cell chirp puncta co-detected by xAM and puncta detected only by cell chirp. Co-detected puncta had higher signal-to-background ratios and stronger harmonic content: median signal-to-background ratios were 49.5 versus 45.5 dB for HEK-293T cells and 49.5 versus 46.1 dB for MDA-MB-231 cells; harmonic-to-fundamental ratios were −0.91 versus −3.27 dB and −0.73 versus −2.59 dB, respectively. Sample sizes were 299 co-detected and 762 cell chirp-only HEK-293T puncta and 514 co-detected and 892 cell chirp-only MDA-MB-231 puncta. Punctum distributions are descriptive; sequence and co-detection comparisons used wells as the independent units (n = 5 per cell line). A detection required at least three contiguous axial samples above threshold. Phantom volumes were acquired at 100-µm elevational steps, and matching detections in adjacent planes were consolidated before counting (Methods).

**Extended Data Fig. 5.**
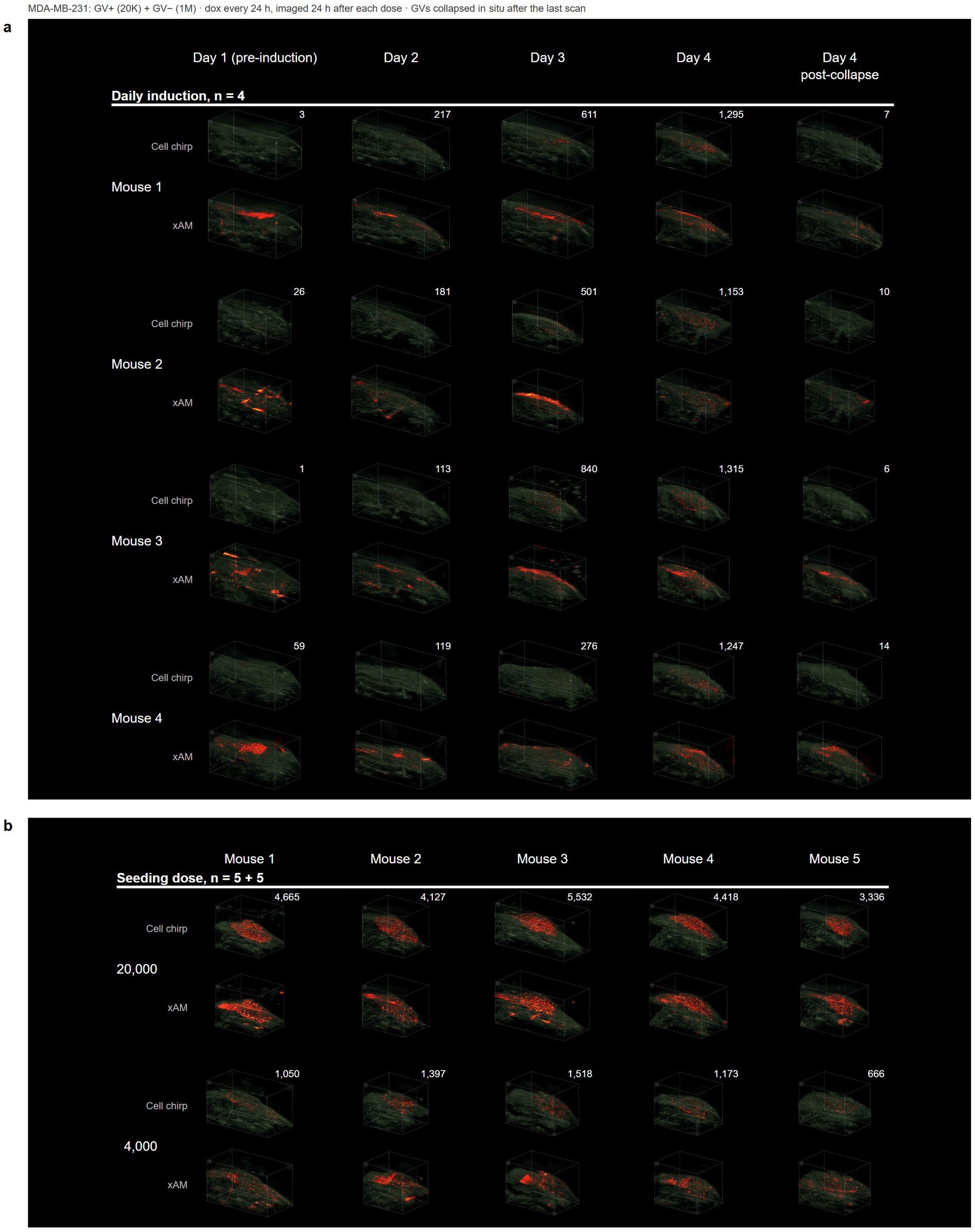
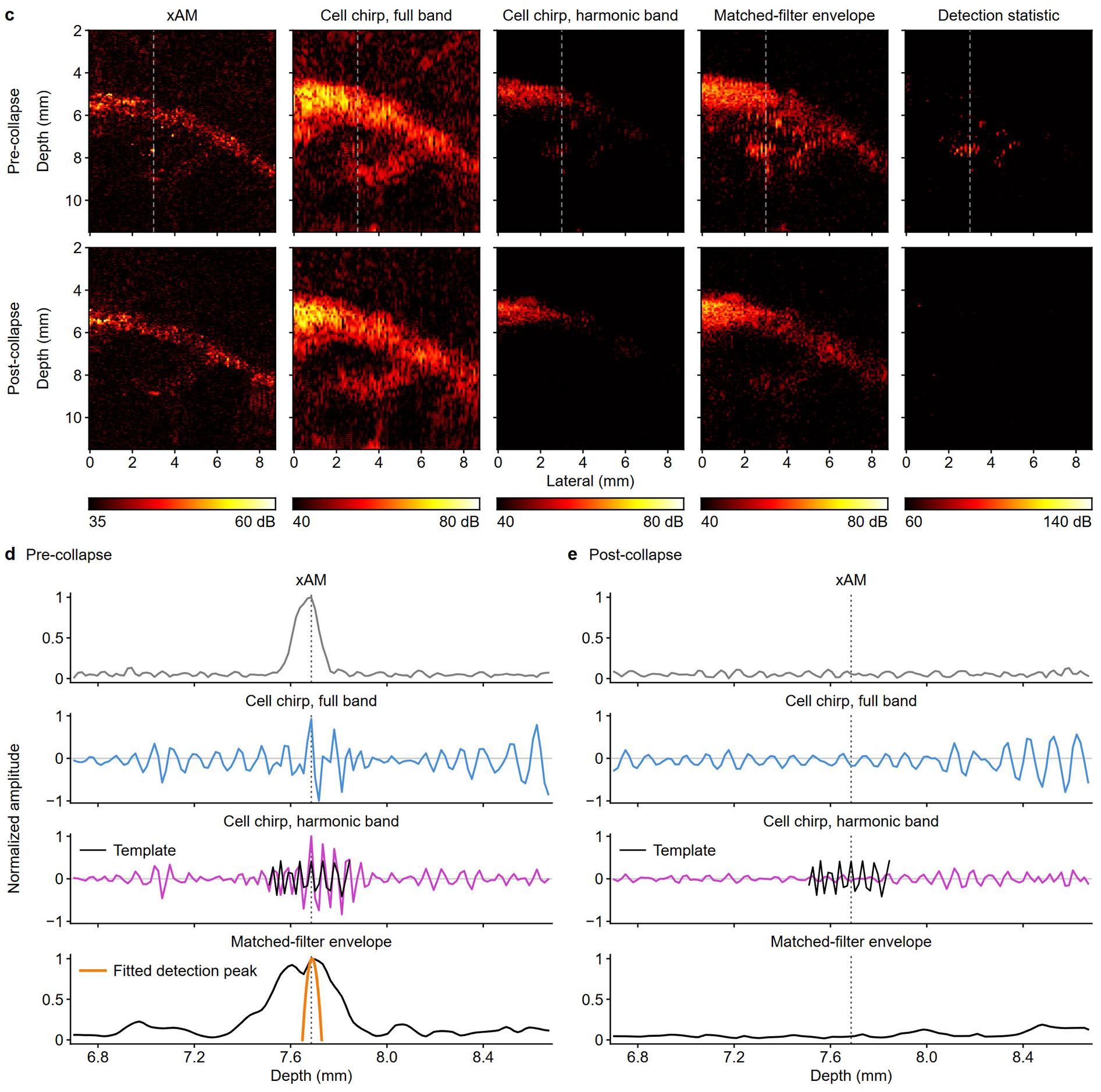
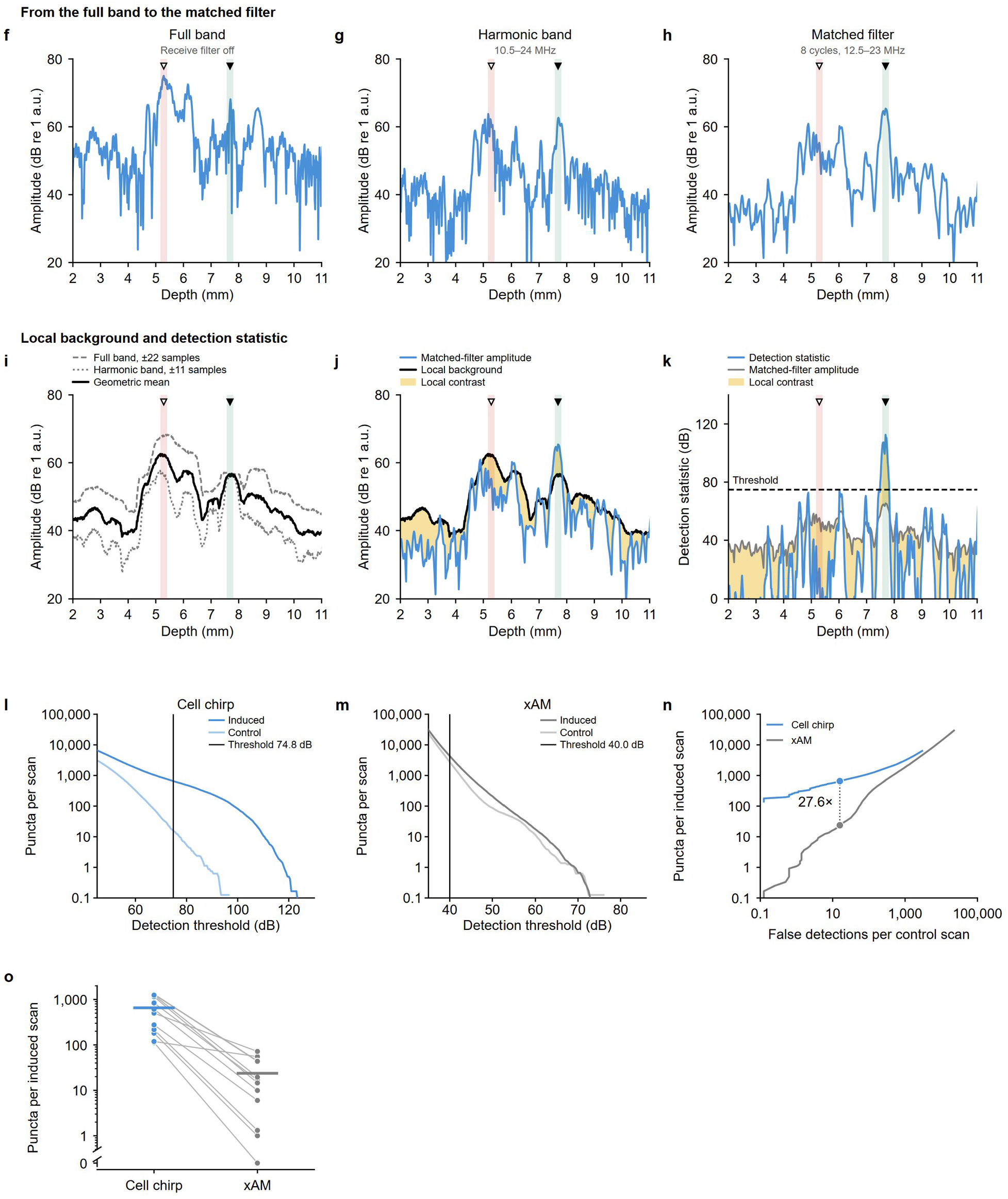
Complete in vivo datasets show induction-dependent reporter recovery, sensitivity to reporter-cell input and GV-dependent signal recovery. a, Complete paired induction dataset from four tumors initiated with 2 × 10⁴ reporter-positive MDA-MB-231 cells among 1 × 10⁶ reporter-negative cells. Each mouse occupies a pair of cell chirp and xAM rows. Columns show the pre-induction scan, three scans acquired 24 h after successive daily doxycycline doses and the post-collapse scan acquired after the final induced measurement. Cell chirp detections increased after the first dose and through subsequent daily measurements in every tumor, then returned near the pre-induction level after GV collapse. Numbers above cell chirp volumes are absolute detection counts. b, Complete reporter-cell input experiment. Separate tumors were initiated with either 20,000 or 4,000 reporter cells among 1 × 10⁶ reporter-negative cells and imaged on day 7 (n = 5 tumors per group). Each column denotes one animal, with paired cell chirp and xAM rows. Cell chirp recovered reporter-associated puncta in every tumor initiated with the 0.4% reporter-cell population and resolved the lower reporter-cell input. All renderings use the same physical scale and crop; cubes indicate 0.5 mm. xAM volumes are included as the non-destructive comparator and are displayed without count overlays. The representative volumes in Fig. 4b are drawn from these complete datasets. c–e, In vivo verification of cell chirp signal recovery in an induced reporter-expressing tumor before and after GV collapse. c, Images before collapse (top) and after collapse (bottom), shown at the same color limits. Columns show the xAM envelope, full-band cell chirp image, 10.5–24-MHz harmonic-band image, matched-filter envelope and final detection statistic. The dashed line marks the raylines analyzed in d and e. Detections required at least three contiguous axial samples above the fixed 74.8-dB in vivo threshold. d, Before collapse, the reporter-associated echo retained the chirp code and produced a compressed detection peak. e, After collapse, this peak disappeared while full-band tissue echoes persisted. The rayline was selected from the induction dataset in a. f–k, Measured version of the detection sequence in Fig. 2b**–g****, along** the pre-collapse rayline in c (the same punctum), processed with the settings used for every in vivo count (Methods). The green band and filled triangle mark the reporter-associated punctum; the red band and open triangle mark a bright tissue echo. f, Full-band envelope with the receive filter disabled. g, Harmonic-band envelope after 10.5–24-MHz filtering. h, Matched-filter amplitude obtained with the eight-cycle, 12.5–23-MHz template. i, Local background estimates from the full-band (±22 samples) and harmonic-band (±11 samples) envelopes, with the central response excluded, and their geometric mean. j, Matched-filter amplitude relative to the local background; shading denotes local contrast. k, Detection statistic with the fixed 74.8-dB in vivo threshold; only samples at the punctum cross it. Panels f–j share a 20–80-dB amplitude scale referenced to 1 arbitrary unit. l,m, In vivo threshold sweeps for cell chirp (l) and xAM (m) in the four-animal induction cohort. Induced scans comprised three post-induction time points per animal (n = 12 scans); controls comprised one pre-induction and one post-collapse scan per animal (n = 8 scans). Vertical lines mark the fixed detection thresholds (74.8 dB for cell chirp and 40.0 dB for xAM). n, Mean puncta per induced scan against mean false detections per control scan as the detection threshold is swept, from the scans in l and m. Points mark cell chirp at its fixed threshold (15.75 false detections per control scan and 655.7 puncta per induced scan) and xAM at the threshold giving the same false-detection rate (23.72 puncta per induced scan), interpolated between adjacent swept thresholds; cell chirp detected 27.6 times as many puncta. o, Puncta in each of the 12 induced scans at these two thresholds. Lines join the two sequences in the same scan and bars mark the means. One scan had no xAM detection and is plotted at 0, below the axis break. For l–o, a detection required at least three contiguous axial samples above threshold. In vivo volumes were acquired at 200-µm elevational steps, and detections were counted separately in each plane and summed without cross-plane consolidation (Methods).

**Extended Data Fig. 6.**
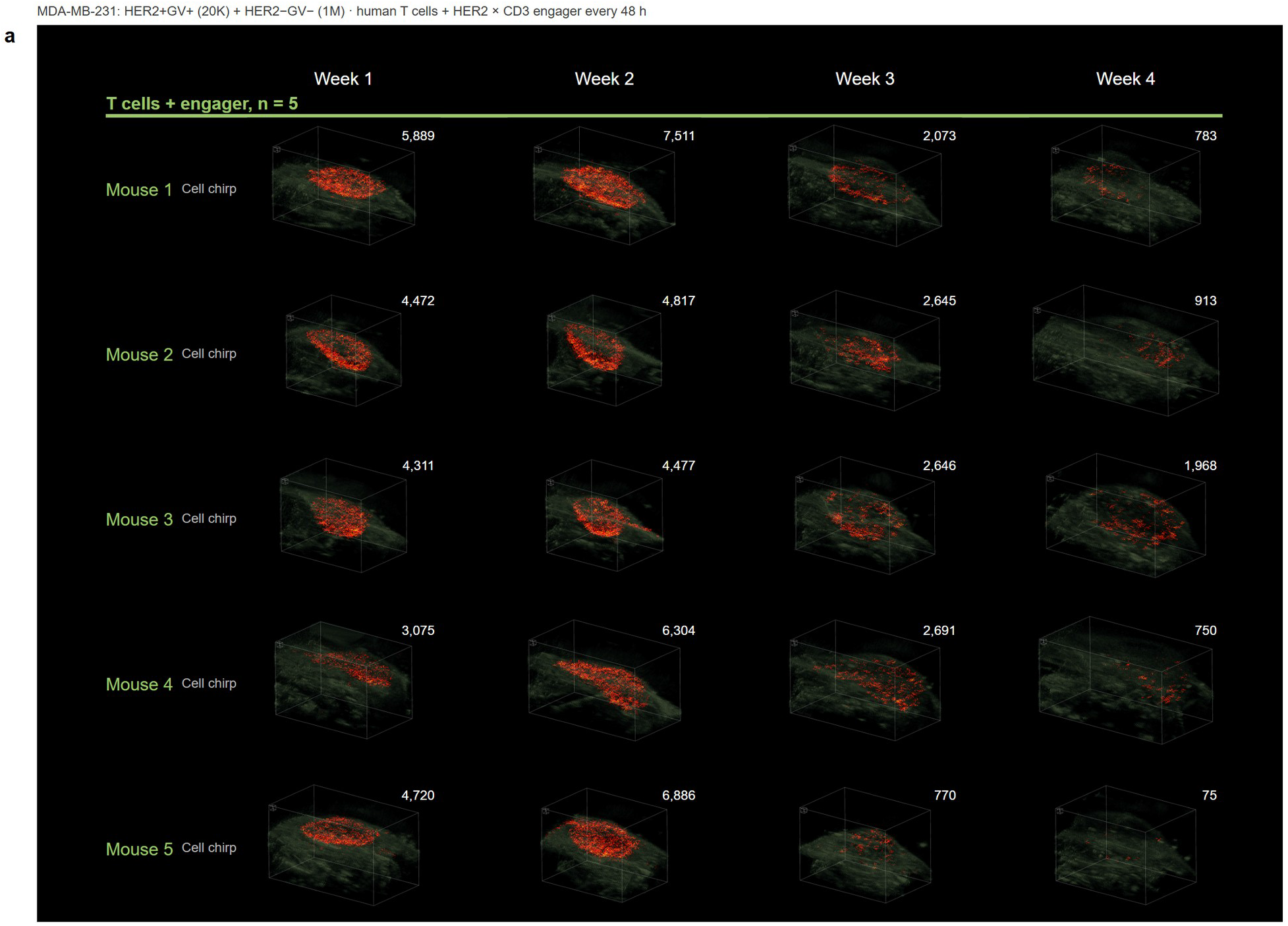
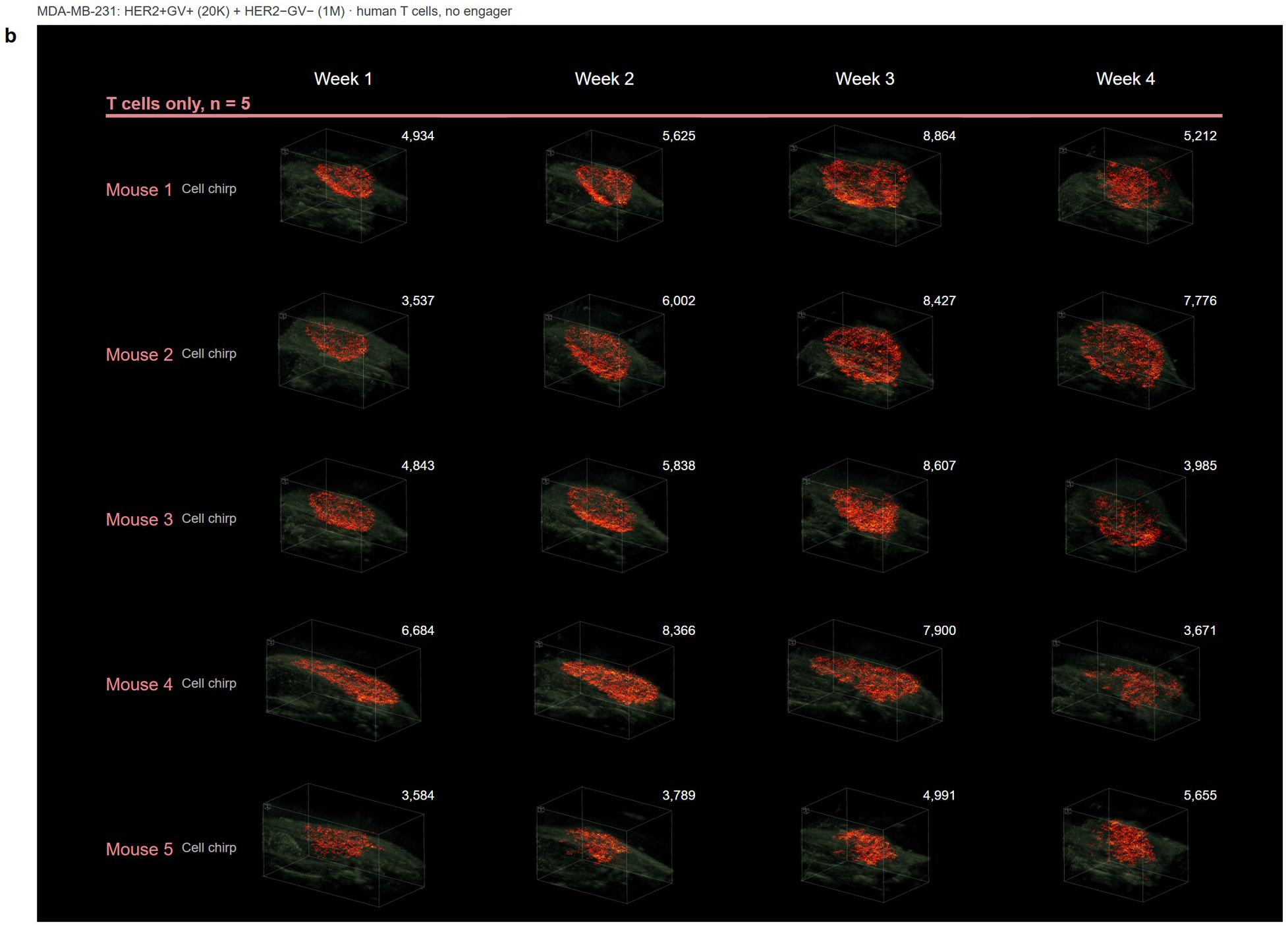
Complete longitudinal dataset for the HER2-positive reporter population. a,b, Complete longitudinal dataset for the antigen-targeted reporter experiment. Tumors contained 2 × 10⁴ reporter-positive, HER2-positive MDA-MB-231 cells among 1 × 10⁶ reporter-negative parental cells without ectopic HER2 expression. All mice received human T cells after week 2. Mice in a also received a HER2×CD3 BiTE every 48 h; mice in b received T cells alone (n = 5 mice per group). Weeks 1 and 2 preceded treatment, and weeks 3 and 4 followed treatment. Each mouse occupies one row of cell chirp volumes, with weekly scans arranged in columns. All ten mice and all four time points are shown. Numbers above the volumes are absolute detection counts. All renderings use the same physical scale and crop; cubes indicate 0.5 mm. Reporter detections declined after treatment in the T cells + BiTE group and remained higher in T-cell-only controls. Mouse 1 in each group is shown in Fig. 5b.

**Extended Data Fig. 7.**
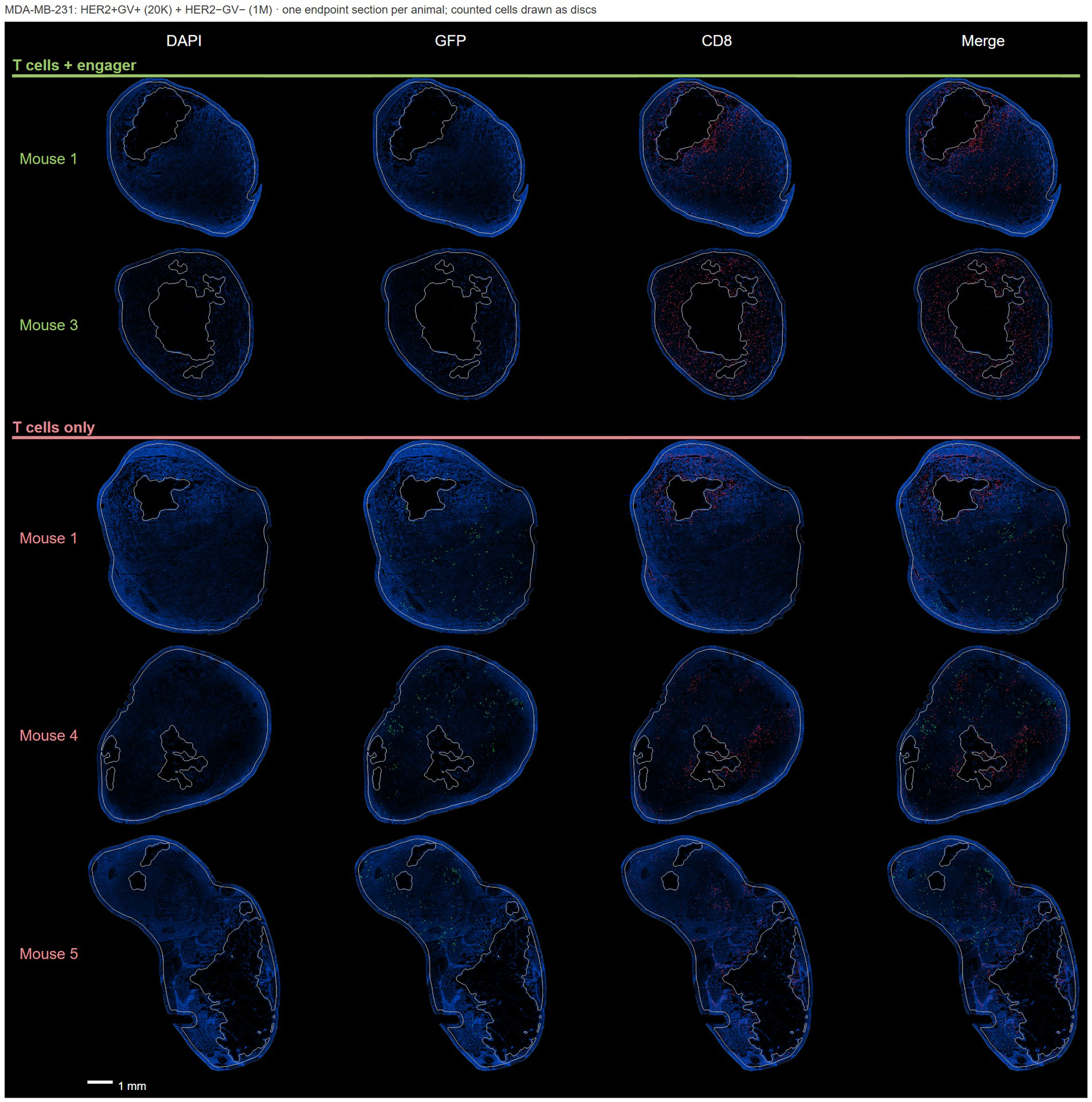
Endpoint microscopy in the HER2-positive reporter cohort. Complete endpoint confocal dataset used to evaluate the antigen-targeted reporter population. Tumors contained 2 × 10⁴ reporter-positive, HER2-positive cells among 1 × 10⁶ reporter-negative parental cells without ectopic HER2 expression. Sections are shown from the two analyzed mice that received T cells + BiTE and the three analyzed mice that received T cells alone. Columns show DAPI, GFP, CD8 and merged channels. Colored discs mark counted GFP-positive or CD8-positive cells; outlines denote the tumor and excluded internal regions. BiTE-treated tumors contained fewer GFP-positive reporter cells and more CD8-positive cells, corroborating the longitudinal acoustic response in Fig. 5. Scale bar, 1 mm.

**Extended Data Fig. 8.**
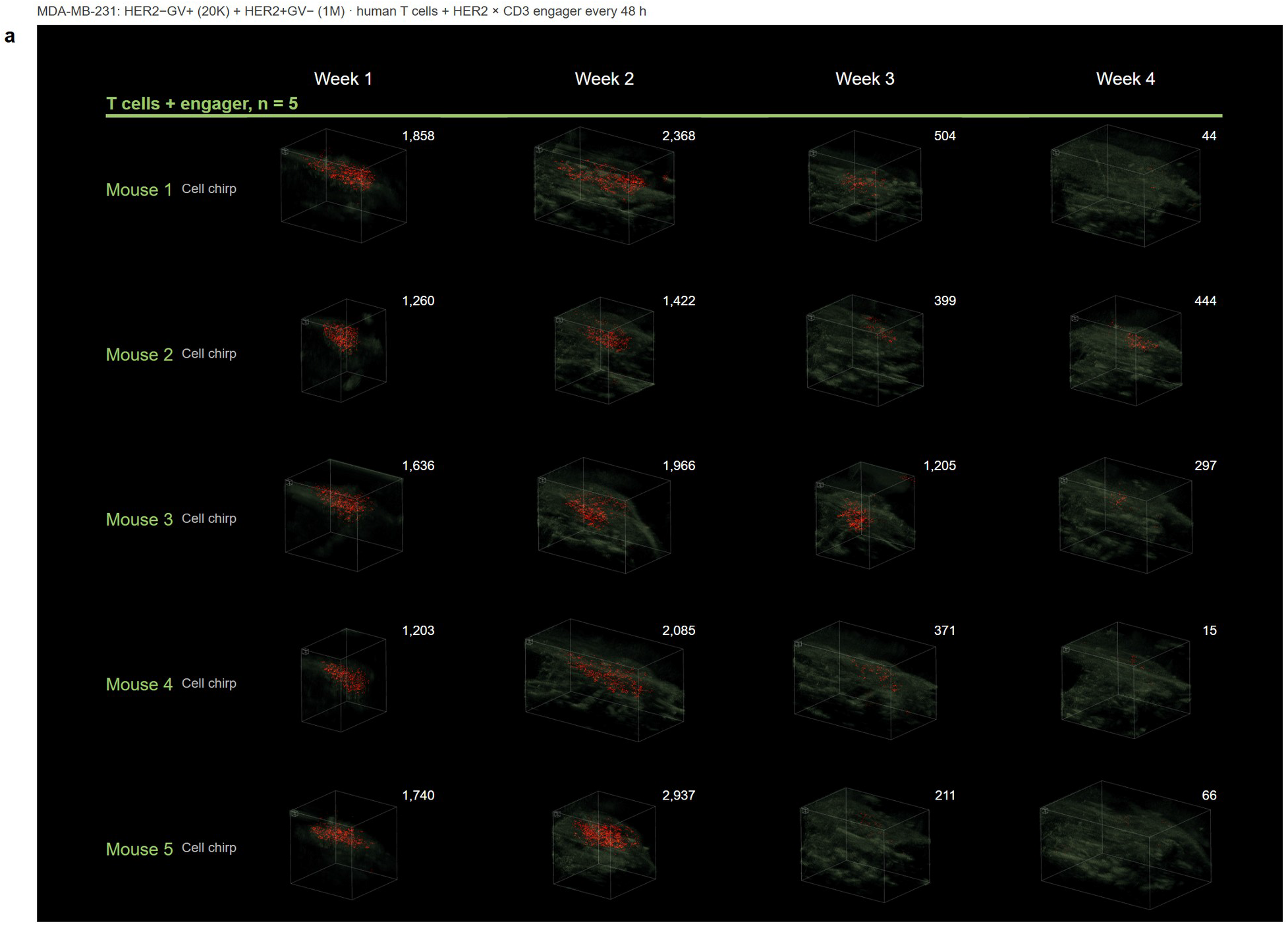
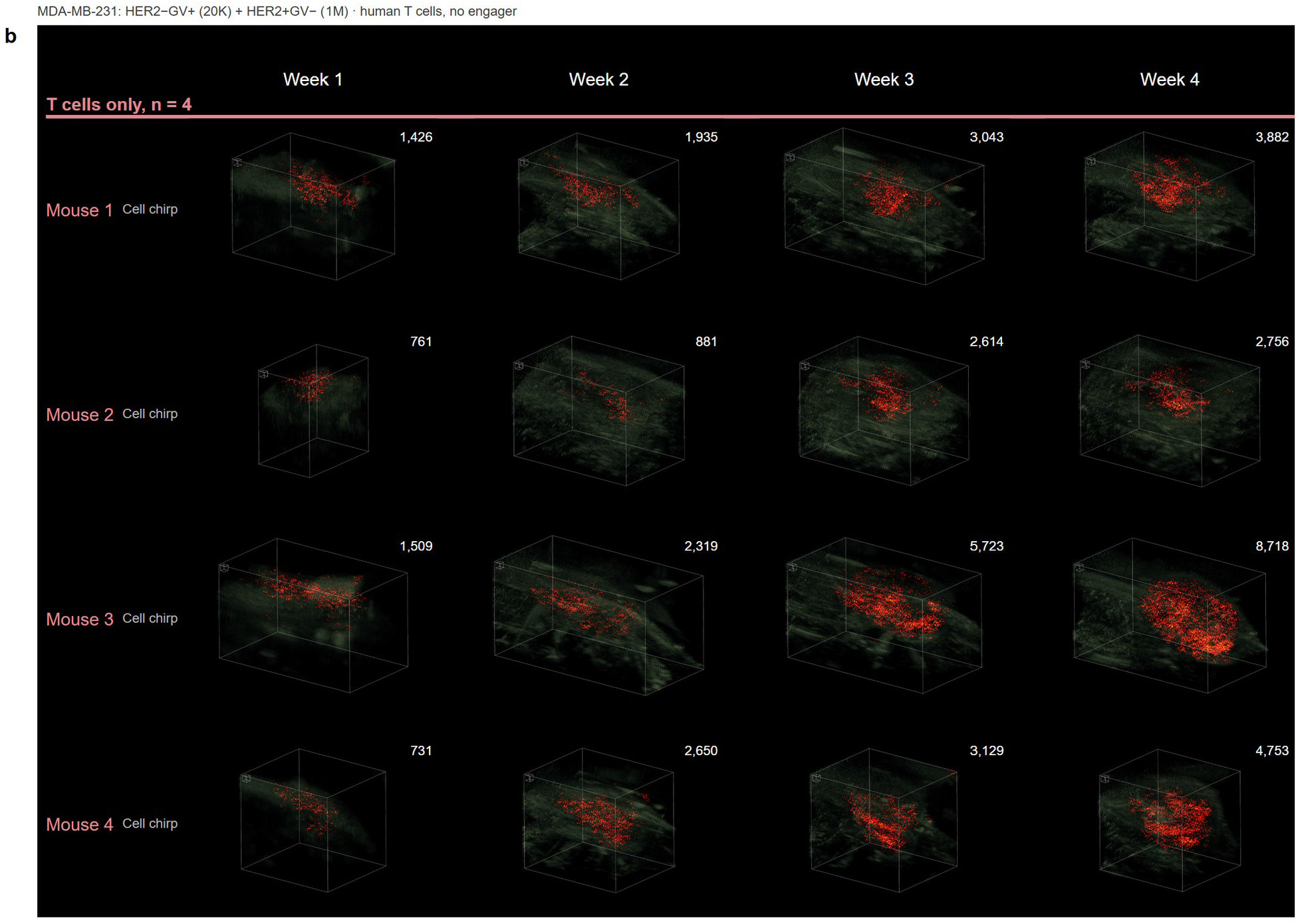
Complete longitudinal dataset for the HER2-negative reporter population. a,b, Complete longitudinal dataset for the antigen-negative reporter experiment. Tumors contained 2 × 10⁴ reporter-positive, HER2-negative MDA-MB-231 cells among 1 × 10⁶ HER2-positive, reporter-negative cells, placing the reporter in cells not recognized by the BiTE. All mice received human T cells after week 2. Mice in a also received the HER2×CD3 BiTE every 48 h (n = 5); mice in b received T cells alone (n = 4). Weeks 1 and 2 preceded treatment, and weeks 3 and 4 followed treatment. Each mouse occupies one row of cell chirp volumes, with weekly scans arranged in columns. All nine mice and all four time points are shown. Numbers above the volumes are absolute detection counts. All renderings use the same physical scale and crop; cubes indicate 0.5 mm. Reporter detections declined after treatment in every T cells + BiTE tumor and increased in every T-cell-only control. Mouse 1 in each group is shown in Fig. 6b.

**Extended Data Fig. 9.**
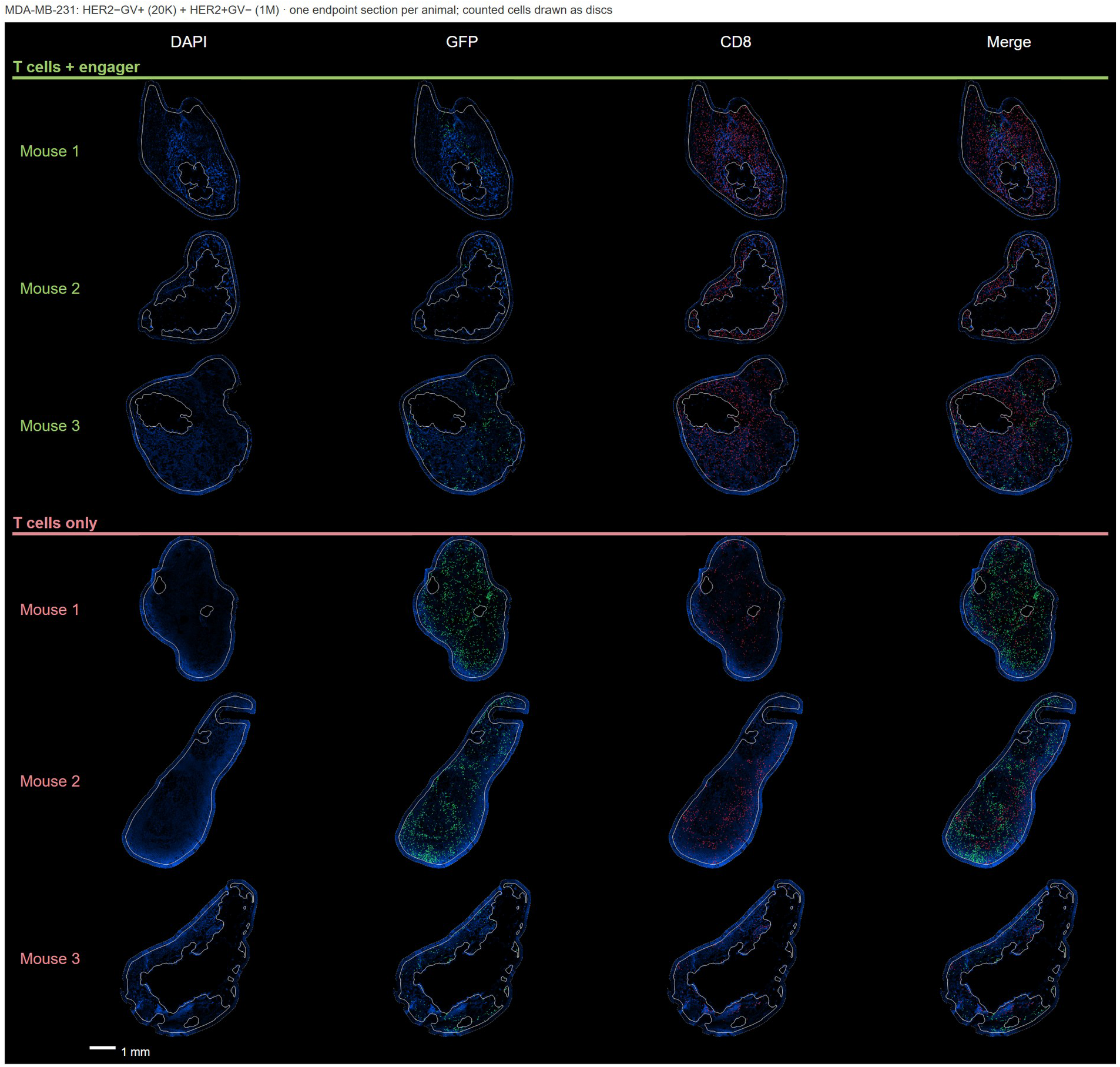
Endpoint microscopy in the HER2-negative reporter cohort. Complete endpoint confocal dataset used to evaluate the antigen-negative reporter population. Tumors contained 2 × 10⁴ reporter-positive, HER2-negative cells among 1 × 10⁶ HER2-positive, reporter-negative cells. Sections are shown from three mice that received T cells + BiTE and three that received T cells alone. Columns show DAPI, GFP, CD8 and merged channels. Colored discs mark counted GFP-positive or CD8-positive cells; outlines denote the tumor and excluded internal regions. BiTE-treated tumors contained fewer GFP-positive reporter cells and a higher density of CD8-positive cells, corroborating the longitudinal response in Fig. 6. Scale bar, 1 mm.

